# A pathway to next-generation mast cell stabilizers identified through the novel Phytomedical Analytics for Research Optimization at Scale data platform

**DOI:** 10.1101/2025.08.30.673232

**Authors:** C. Jansen, B.G. Rice, B. Wooton, J. Howard, S. Elmasri, T. Rivera, L.M.N. Shimoda, A.J. Stokes, C.N. Adra, A.L. Small-Howard, H. Turner

## Abstract

Mast cell stabilizers (MCS) have the potential to address unmet therapeutic need in allergy and inflammation management. MCS chronically suppress all arms of the pro-inflammatory mast cell response to stimulation (e.g., histamine, protease, lipid mediators, cytokines, chemokines). They may therefore outperform approaches such as H1, H2 and H4 inhibitors (antihistamines) which block only the acute histamine release by mast cells and leave the rest of the functional response untouched. Despite their potential, current MCS (e.g., cromolyn sodium, Tranilast, nedocromil) in clinical use are hindered by poor bioavailability, frequent dosing, long lags to onset of relief, and enigmatic mechanisms of action. MCS have their origins in phytomedicine: cromolyn sodium is the longest standing drug in the class and is a derivatized form of Khellin from *Ammi visnaga,* used as an anti-inflammatory. Other phytopharmacopeias may offer candidate ‘next generation’ MCS (ngMCS) and in this study we hypothesized that a coupled pharmacoanalytic and *in vitro* pharmacology approach could be used to identify, prioritize and derisk additional candidate MCS from phytomedical sources for later pre-clinical and clinical evaluation. Here, we report a novel data analytics workflow starting with a newly developed phytopharmacopeia data platform with >3.5B linkage pathways (country → medical system → formulation → indication → ingredient organism → chemical component → other parameters), covering 22 M sq. miles of biogeography and historical and contemporary timeframes. Additional data layers include druggability indices, target and pathway analyses. The current study validates a subset of candidate phytomedical ngMCS using *in silico* workflow and *in vitro* pharmacology, and develops a new harmonic mean-based ‘MCS score’ for further streamlining of the candidate prioritization process. This proof-of-concept study may have particular relevance for complex presentations such as Mast Cell Activation Syndrome (MCAS) where ngMCS may outperform current management approaches.

## Introduction

Mast cells drive hypersensitive responses to typically harmless substances. The allergic response initiates when an allergen interacts with polyvalent IgE-FcεRI complexes on the surface of sensitized mast cells, leading to receptor aggregation [1]. This triggers a complex signaling cascade involving kinase pathways and calcium influx which drive the release of granule-bound preformed mediators (e.g., histamine, chymase and tryptase [2]) as well as the *de novo* synthesis and release of lipid mediators (e.g., prostaglandins and leukotrienes), cytokines and chemokines [3]. Mast cells are important therapeutic targets for conditions such as asthma, atopy, urticaria, allergic rhinitis, and allergic conjunctivitis. Currently, several therapeutic approaches target mast cells, with distinct mechanisms of action, advantages, and limitations. *Antihistamines* are common, relatively inexpensive, histamine H1 receptors blockers. They reduce itching, sneezing, and hives, but they do not prevent release of other mediators, and their side effect profiles include drowsiness and dry mouth, impacting patient compliance. *Corticosteroids* administered systemically or topically suppress immune responses including mast cell activation. They are effective, but long-term use is associated with broad immunosuppression, osteoporosis, and adrenal suppression. Desensitization means that quadruple dose escalations are not unusual [4,5]. *Biologics*, (e.g., IgE mAb omalizumab) target severe allergic responses effectively [6] but are expensive and may require parenteral administration, limiting accessibility and convenience. Finally, *mast cell stabilizers* (MCS) prevent both degranulation of mast cells and release of multiple inflammatory mediators at the point of origin, offering a potentially comprehensive therapeutic approach and prophylaxis. Unlike antihistamines, which primarily block histamine receptors and act after mast cell mediator release has occurred, MCS seem to target underlying pathways that control multiple arms of the MC mediator release responses. MCS potentially offer a more complete approach to managing allergic responses through blocking more than one ‘arm’ of the mast cell response to antigen/allergen stimulation [7–9]. Temporally, MCS effects are chronic and less subject to desensitization and hence dose-escalation than antihistamines and corticosteroids. This multi-dimensional suppression of MC-mediated tissue inflammation may be particularly important in long-term allergy management and in the recently-recognized syndromic presentation of MCAS (Mast Cell Activation Syndrome) [10–13] which cannot be managed by anti-histamines and corticosteroids alone [14]. MCS that are in current clinical use with several decades of therapeutic history include cromolyn sodium, nedocromil, ketotifen and Tranilast. Their clinical utility is limited by low bioavailability, the need for frequent dosing, and some problematic side-effect profiles [15,16]. Several are limited to ocular indications, and Tranilast [17] has recently been sidelined due to side effect issues. In concert with the acute relief provided by antihistamines, MCS offer a potentially important therapeutic option but there is a need to identify next-generation MCS (ngMCS) with increased potency, longer duration of action, decreased costs, improved patient adherence through less frequent dosing and for which a mechanism of action is fully understood.

MCS originated in a phytomedical space, with Middle Eastern, African and Indian subcontinent medicinal systems leveraging *Ammi visnaga* (Khella) seeds for treatment of allergy and asthma [18]. Western rediscovery of this therapeutic approach occurred in the 1960s when Altounyan [19] isolated an active factor (Khellin) which was later derivatized to cromolyn sodium, increasing its ionic character and improving pharmacokinetics and bioavailability. In 2013 Finn and Walsh [20] performed a metareview of the current unmet therapeutic need for ngMCS and identified 27, predominantly phytochemical, natural products that met certain criteria for classification as potential MCS, including documented efficacy in chronic passive cutaneous anaphylaxis assays *in vivo*. They identified a diverse and heterogeneous phytochemical candidate set including flavonoids [21–30], coumarins [31–36], phenols [37–39], terpenoids [40–43] and amino acids [44] on the basis of literature review. However, there has been no strategic follow up to this study and we continue to see work positing the need for strategic development of ngMCS.

Conventional drug discovery pipelines take ∼15 years, cost $2B [4] and see significant (∼45%) attrition [4]. Phytomedicines used to treat MC disorders in global medicine systems (GMS) are attractive starting points for development. However, their complex polypharmacy (formulations are typically tens of organisms and hundreds to thousands of primary and secondary metabolite compounds) is a barrier to rational design and regulatory pathways. In this study we hypothesize that a coupled pharmacoanalytic and *in vitro* pharmacology approach could be used to identify, prioritize and derisk additional candidate MCS from phytomedical sources for later pre-clinical and clinical evaluation. We tested the application of a new data platform and phytochemical drug discovery platform (PhAROS^TM^, Phytomedicine Analytics for Research Optimization at Scale), initially using the Finn and Walsh ngMCS candidate as a semi-validated test compound set, and progressing to predictive analytics leveraging the entire phytochemical space in PhAROS^TM^. We tested a PhAROS^TM^ workflow called in silico convergence analysis (ISCA) where candidates are raked and prioritized based on their representation in GMS formulations for a specific indication (in this case MC-related inflammatory disorders). We then tested the Finn/Walsh compound set performance using PhAROS^TM^ data layers that examine druggability indices and target analysis to further discern between candidates for advancement in a development pipeline. We developed and tested a novel ‘MCS score’ index that integrates across PhAROS^TM^ data layers on each compound and used it predictively to develop a novel ngMCS candidate list both in terms of chemical class and individual compounds. Using the Finn/Walsh compound set as a ‘ground truthing’ resource provided reciprocal validation both for the PhAROS^TM^ method and for the Finn/Walsh compounds. It also yielded additional information that enabled ranking and discernment within this group. Beyond Finn/Walsh, PhAROS^TM^ generated key predictive data on chemical classes and individual compounds that position future preclinical and clinical work on rational design of phytochemical next-generation mast cell stabilizers as both single and combinatorial drug formulations.

## Materials and Methods

### Construction of the PhAROS^TM^ Data Platform

#### Data Acquisition

Traditional medicine system datasets were pulled via site download options or web data extraction from Encyclopedia of Traditional Chinese Medicine (ETCM) [45], Ethnomedicinal Plant Database (ETM-DB) [46], Indian Medicinal Plants, Phytochemistry And Therapeutics (IMPPAT) [47], South African National Chemical Database (SANCDB), Kampo Medicine Database (KampoDB) [48], Kyoto Encyclopedia of Genes and Genomes (KEGG) [49], Northern African Natural Products Database (NANPDB) [50], Taiwanese Indigenous Plant Database (TIPdb) [51], Northeast Asian Traditional Medicine (TM-MC) [52], Traditional Chinese Medicine Integrative Database (TCMID) [53], Bioactive Phytochemicals Molecular Database (BioPhytMol) [54], Medicinal Plant Server (MedPServer) [55], Korean Traditional Knowledge Portal (KTKP) [56], and Pharmacological Database of Korean Medicinal Plants (PharmDB-K) [57]. Each traditional medicine system dataset was preprocessed to include most of the following features: System of origin (TXM), ingredient combination (Formula), ingredient organism scientific name (Species), compounds associated with each species (Compound), compound identifier (CID), and usage (Indications).

#### Data Enrichment

The traditional medicine system datasets were enriched with plant-phytochemical lists from Dr. Duke’s Phytochemical Database and the compounds were further enriched with Pubchem information to include CID, INCHIKEY, and Classification. Missing compound classifications were filled in using Classyfire [58]. Country codes were matched to appropriate ISO 3166 Codes for mapping. Protein targets were matched to compounds via CHEMBL identifiers and bioassays.

### Data Analysis

Exploratory data analysis was conducted via the following python packages: pandas, numpy, matplotlib, and seaborn. Plotly, biophyton, networkx and chembl_web_client were also applied Python packages used in the frequency analysis. We curated ligand-target binding relationships for MCS compounds from ChEMBL v32 and the BindingDB (July 2024 update), restricting associations to those where the target organism was *Homo sapiens*. Additionally, we utilized the ChEMBL Target Prediction API [59,60] to predict targets based on SMILES for all MCS compounds, filtering results to include only *Homo sapiens* targets classified as “active” with a confidence level of at least 90%. Known and predicted ligand-target associations were merged into a single dataset. The ligand-target data was then transformed into network visualizations using the Python package NetworkX. These networks were further explored and visualized with Gephi software, employing the ForceAtlas2 layout algorithm and a subsequent Label Adjust step to prevent label overlap.

### Indication dictionary construction

The indication dictionary delineates the search terms and truncations used to interrogate PhAROS^TM^ with the aim of identifying data fields linked to certain disorders. For the MCS study described here the search terms were designed to focus results on presentations for which MCS would be indicated therapeutics [13,61,62]. Terms used were allerg*, anaphyla*, asthma, bite, congestion, conjunctivitis, dermatitis, eczema, hives, inflamm*, rash, respiratory, rhinitis, sting. Collectively these terms likely capture immune reactions and conditions mast cell stabilizers are commonly used to treat (Table I).

**Table I.**
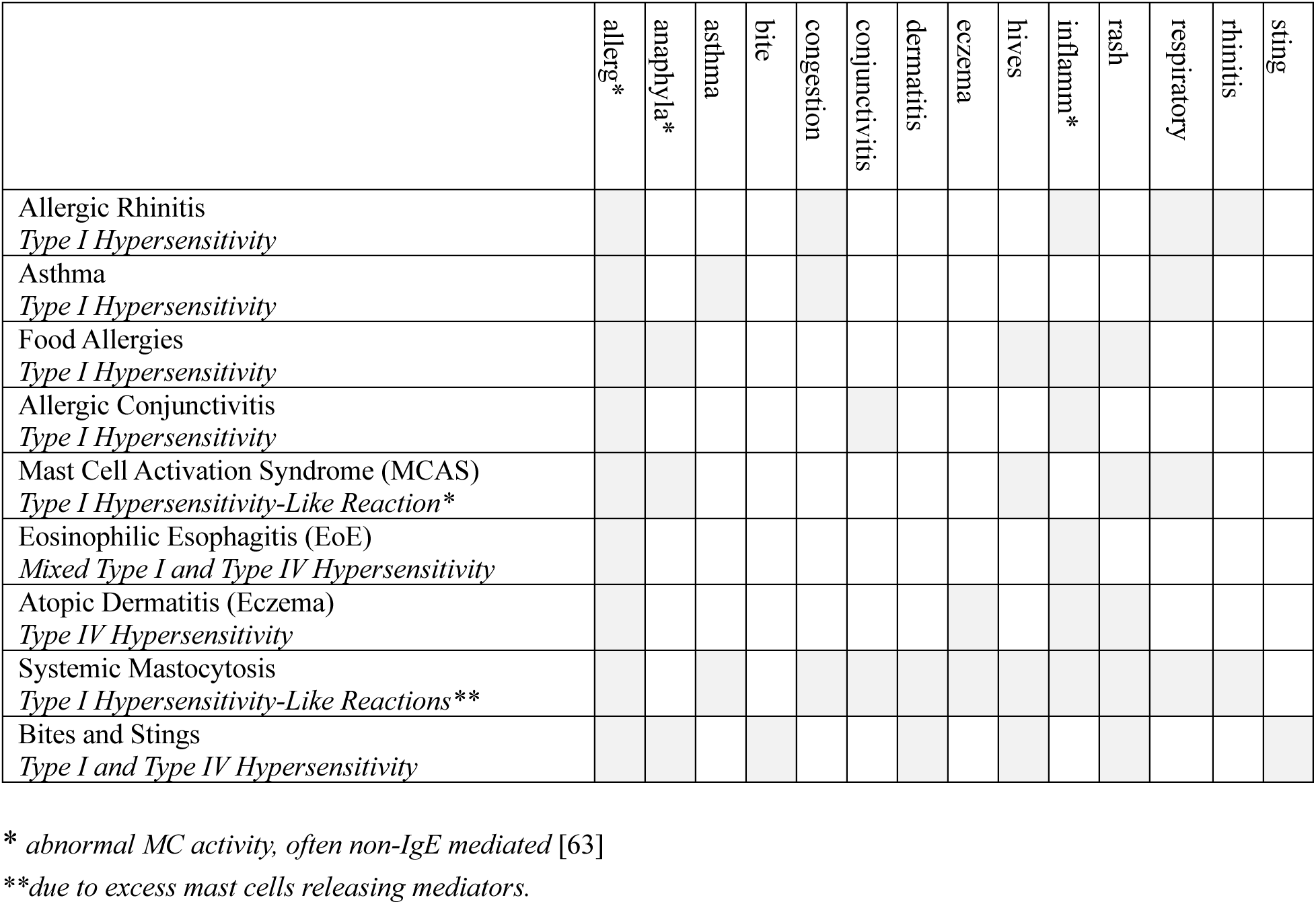
Indication dictionary.

### Beta-hexoseaminidase assay

RBL2H3 cells were cultured in DMEM supplemented with 10% FBS, 5mM Glutamine until 70-80% confluence in a 37°C incubator with 5% CO₂ and 95% humidity. The cells were harvested using Trypsin-EDTA, resuspended in DPBS, and seeded at a density of 1 × 10⁵ to 5 × 10⁵ cells per well in a 96-well microplate. The cells were allowed to adhere for 24 hours. The cells were then stimulated for 30 min with the indicated compound, or via FcεRI (1μg/ml IgE anti-DNP overnight followed by two washes in media then 250ng/ml KLH-DNP for 30-60 minutes at 37°C), with a control group of unstimulated cells included and one set of replicates exposed to 0.05% Tx-100. After stimulation, the supernatant was collected from each well and transferred to a fresh plate. An equal volume of substrate solution (4 mM 4-nitrophenyl-N-acetyl-β-D-glucosaminide (pNAG) in 0.1 M citrate buffer, pH 4.5) was added to both the supernatant and lysate wells and plates were read at 405nm absorbance with A405 correlating with colorigenic beta-hexoseaminidase [64,65] action on the pNAG substrate.

## Results

### Workflow for evaluation of phytomedicine candidates for novel mast cell stabilizers

The PhAROS^TM^ (Phytomedicine Analytics for Research Optimization at Scale) platform integrates formulation-level information, indication dictionaries, multi-source data layers, and curated training sets. It is designed to enable complexity reduction for rational simplification of complex polypharmaceutical phytomedicines. It curates ∼6B potential multiwise linkages across ∼75,000 natural compounds, 50,000 indications and multiple additional data layers. The core workflow includes *in silico* convergence analysis, predictive parameter extraction (e.g., chemical similarity, target overlap, druggability), and novel scoring indices to rank candidate compounds. These are further refined through machine learning and *in vitro* bioassay integration workflows. This approach enables data-driven rational design of single-agent and combinatorial phytochemical drug formulations for inflammatory and allergic indications (Figure 1).

**Figure 1.**
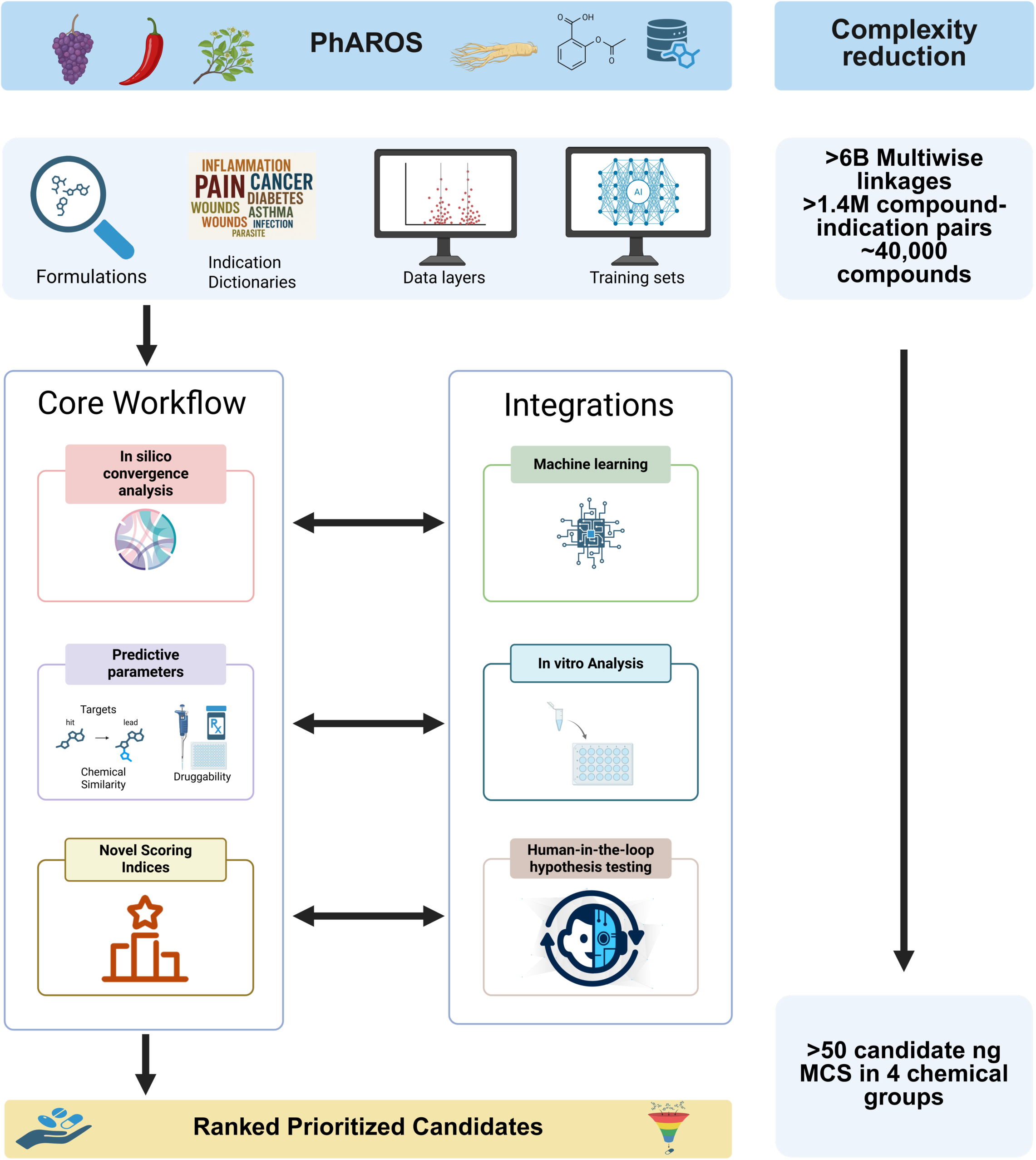
Workflow using PhAROS^TM^ to evaluate potential next generation MCS from phytomedical sources. Schematic overview of the PhAROS^TM^ platform and its core analytic workflow. PhAROS integrates phytomedical data sources including formulations, indication dictionaries, various curated data layers, and training sets. A core analytic workflow for this study begins with in silico convergence analysis, followed by the assessment of predictive parameters such as molecular targets, chemical similarity, and druggability. These analyses inform the development of novel scoring indices used to rank and prioritize compound candidates for further exploration and development. The core workflow is enhanced by two integration modules: machine learning models that refine prediction and classification tasks, and in vitro analysis pipelines that validate pharmacologic activity. The system outputs a ranked list of prioritized natural product-derived multi-compound systems (in this case ngMCS). Created in BioRender (license agreement Jansen, C. (2026) https://BioRender.com/kmobbz6).

### *In silico* convergence analysis of Finn/Walsh candidate ngMCS

Our *in-silico* convergence analysis (ISCA) workflow integrates data from multiple formalized and standardized global phytomedicine systems (GMS) (Figure 2A). We include historical and contemporary Indian (IM), African Medicine (AM), European (EM), Chinese (CM), Japanese (JM), Korean (KM), and South American (SAM) medical systems. In this initial data aggregation effort (Figure 2A), we mined associations between these medicine systems, geographic regions, species, medical indications, formulation, and natural phytochemical compounds across >15 GMS databases and nationally standardized pharmacopeias. This effort was supplemented with mining authoritative text sources in PubMed. We extracted detailed information on natural compounds and medical indications, including compound identifiers (PubChem CID), structural details (InChiKeys, SMILES, IUPAC names), and standardized medical vocabulary (MeSH terms) from sources such as PubChem and KEGG. The outcome is a data platform comprising (Figure 2B, C): 8 traditional medicine systems, over 150 countries, more than 17,000 formulations linked to 50,000 described medical indications, incorporating over 25,000 ingredient organisms and more than 75,000 fully annotated natural compounds. We estimated an upper bound number of potential pairwise linkages at >3B in the data base and multicomponent linkage paths (i.e., country to medical system to formulation to indication to ingredient organism to chemical component plus other data layers such as targets and druggability parameters) at an upper bound of 6B.

**Figure 2.**
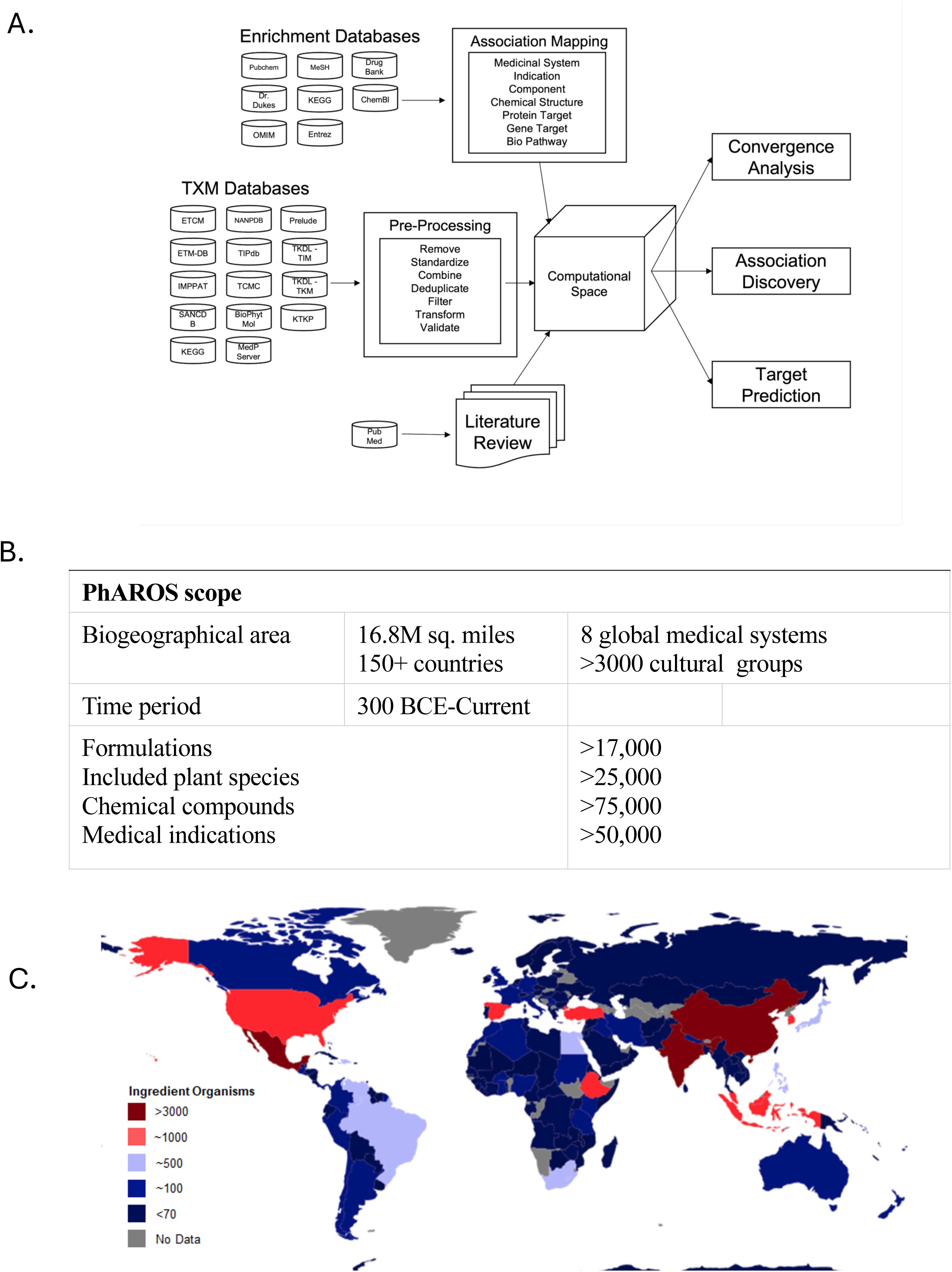
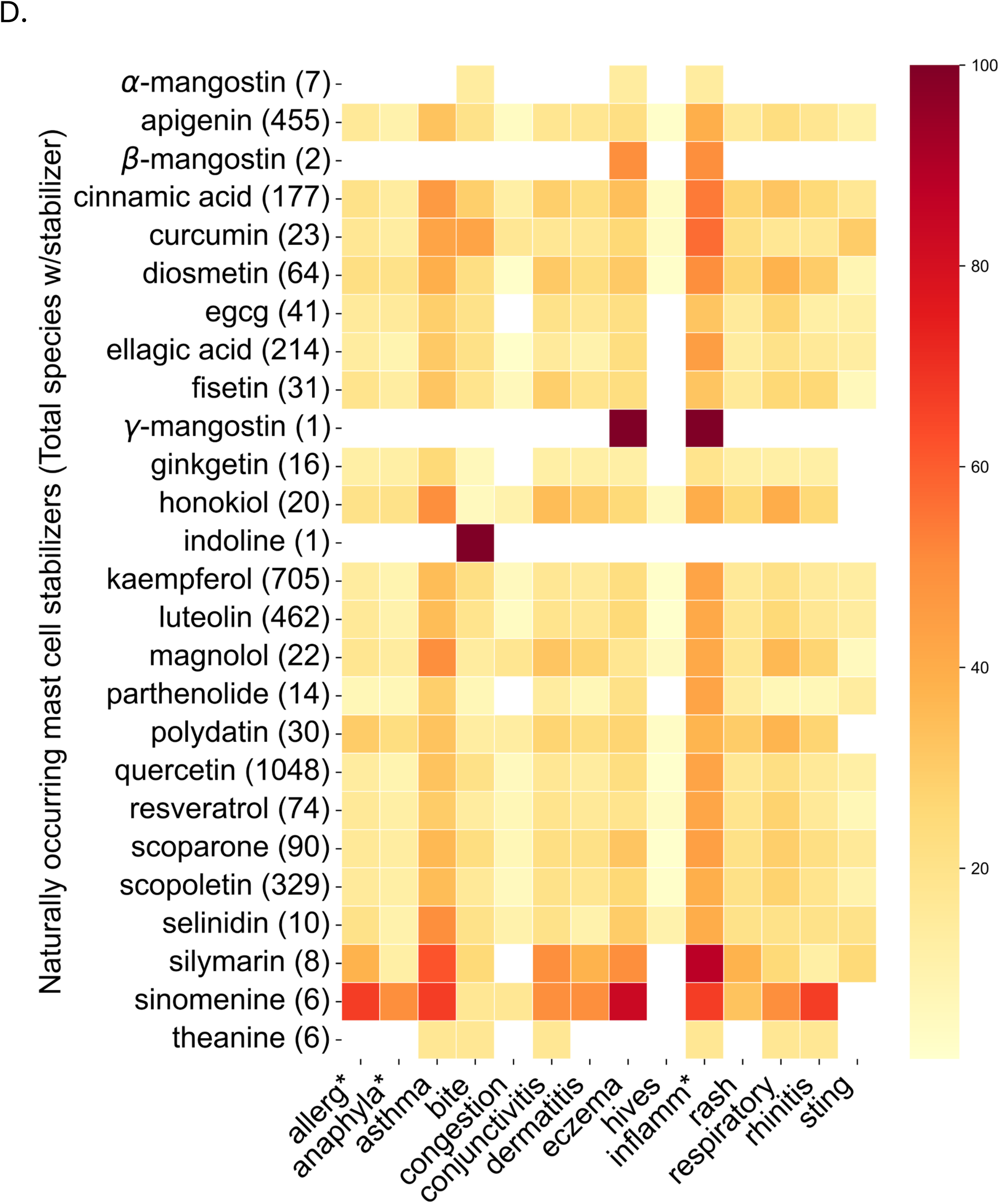
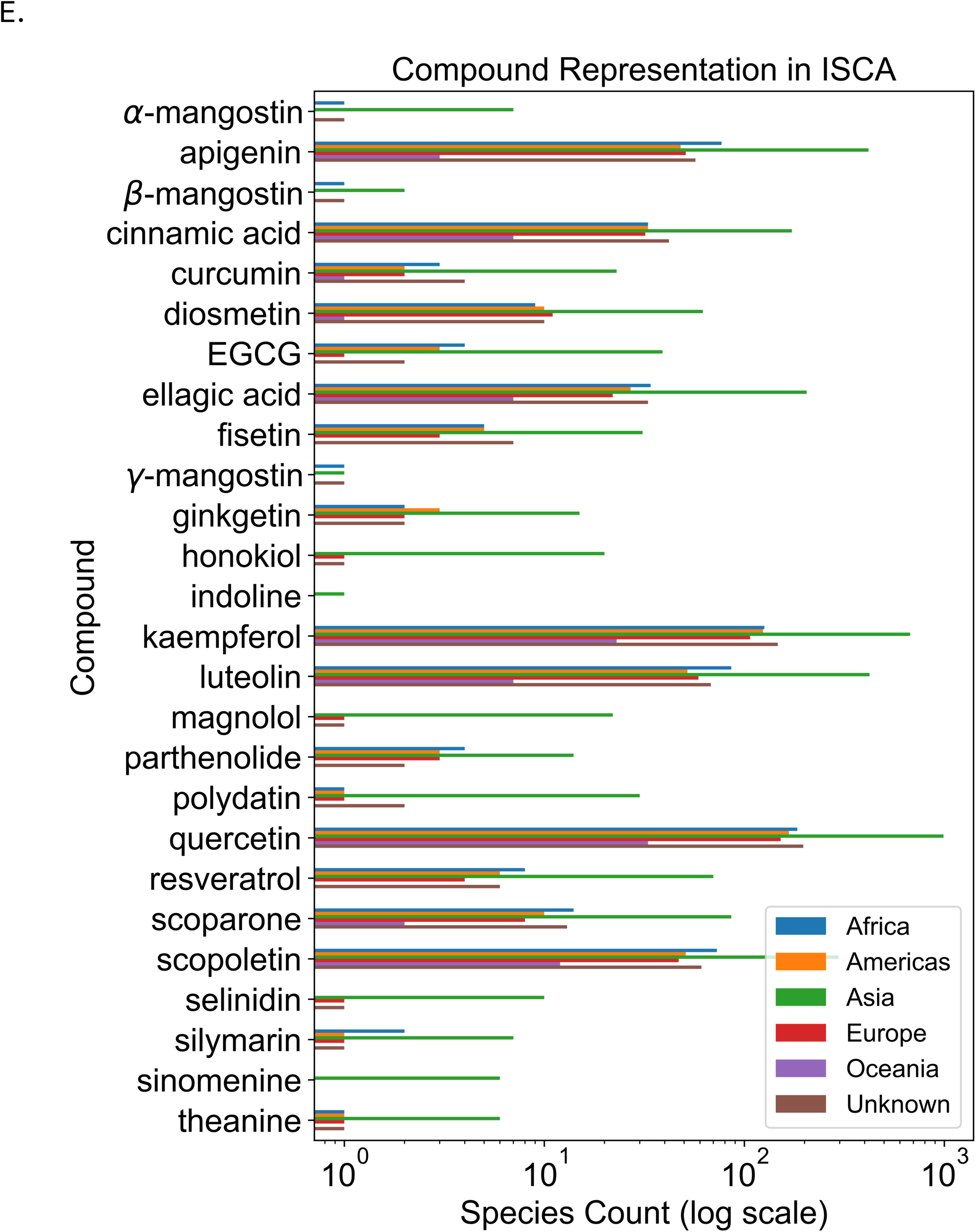
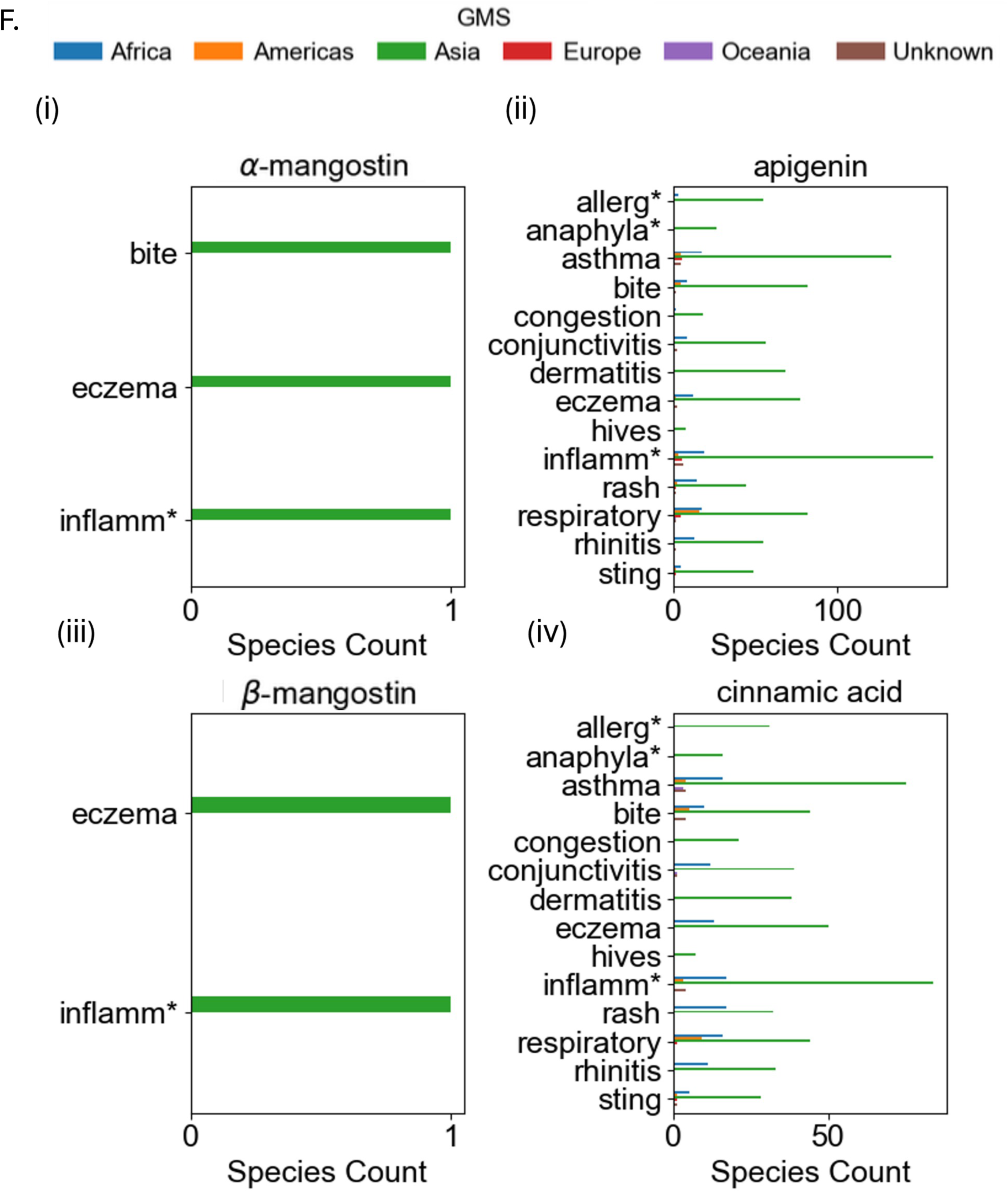
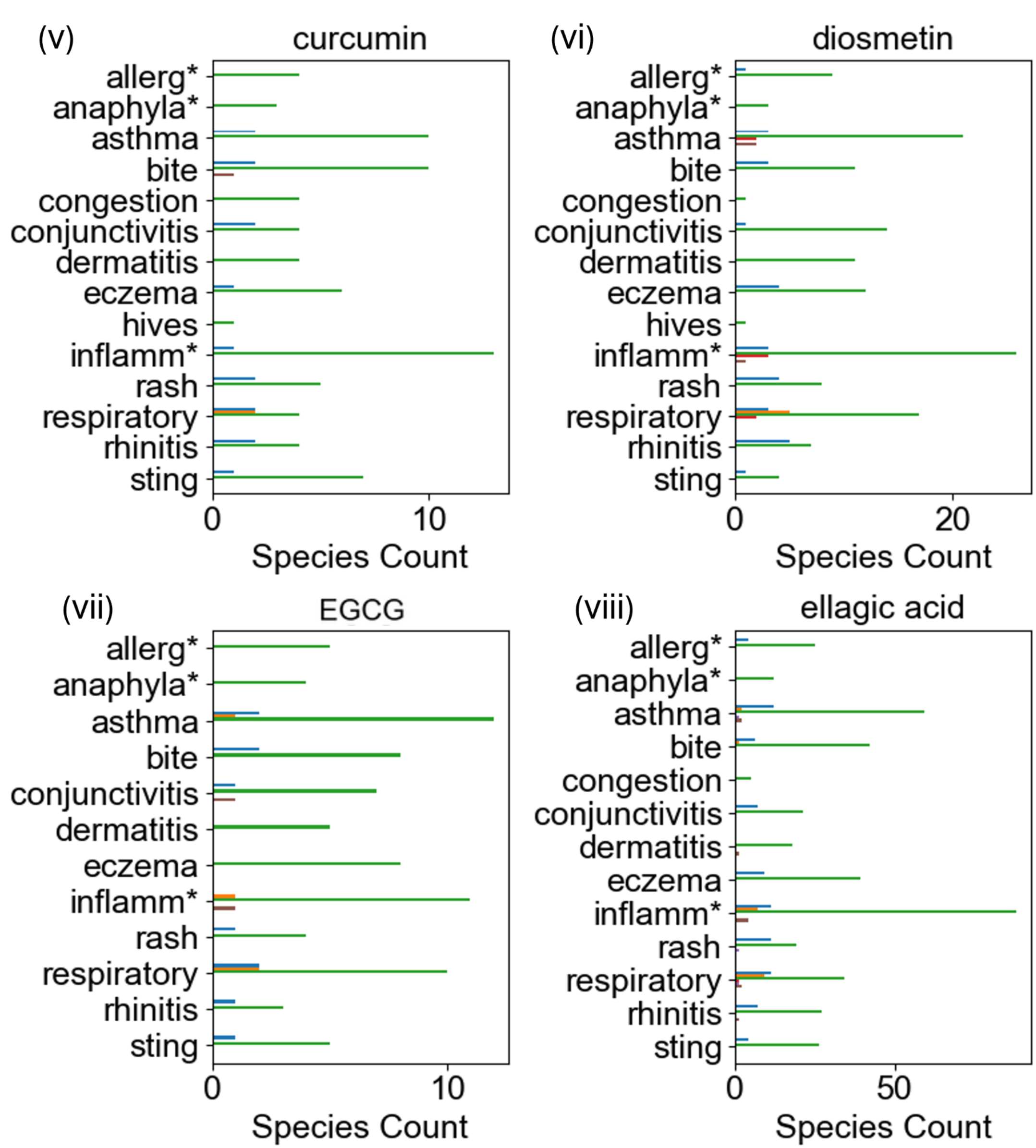
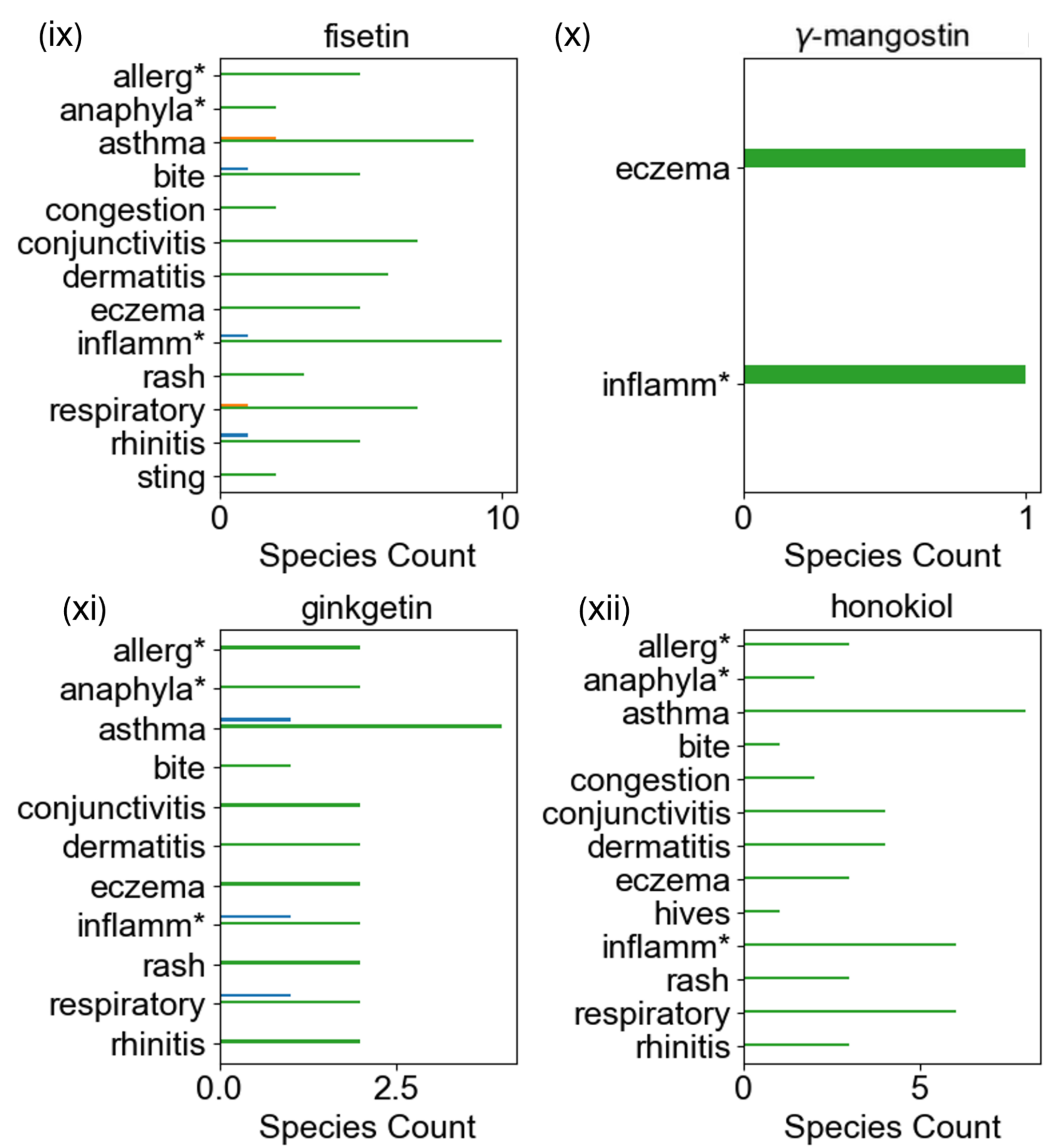
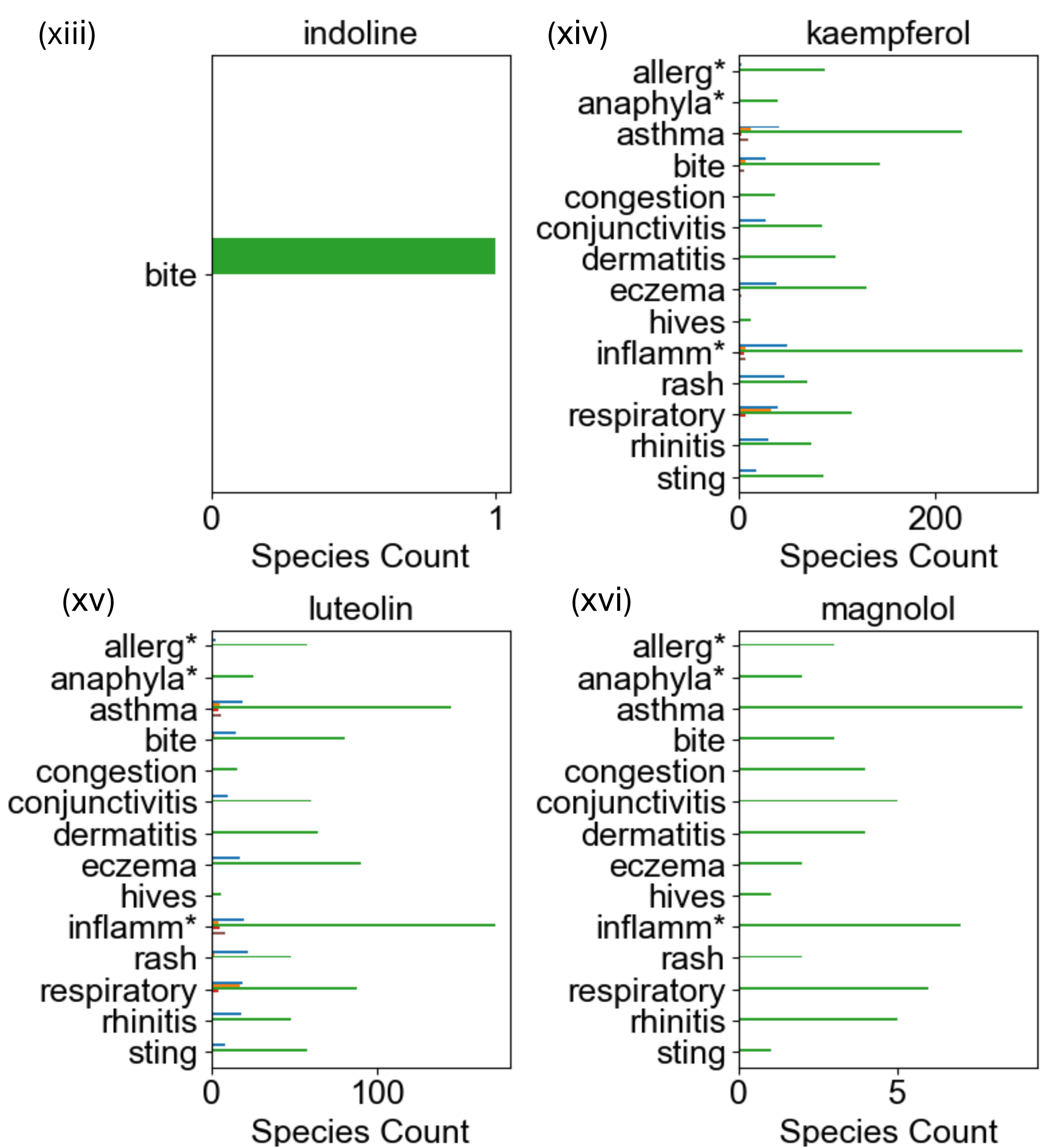
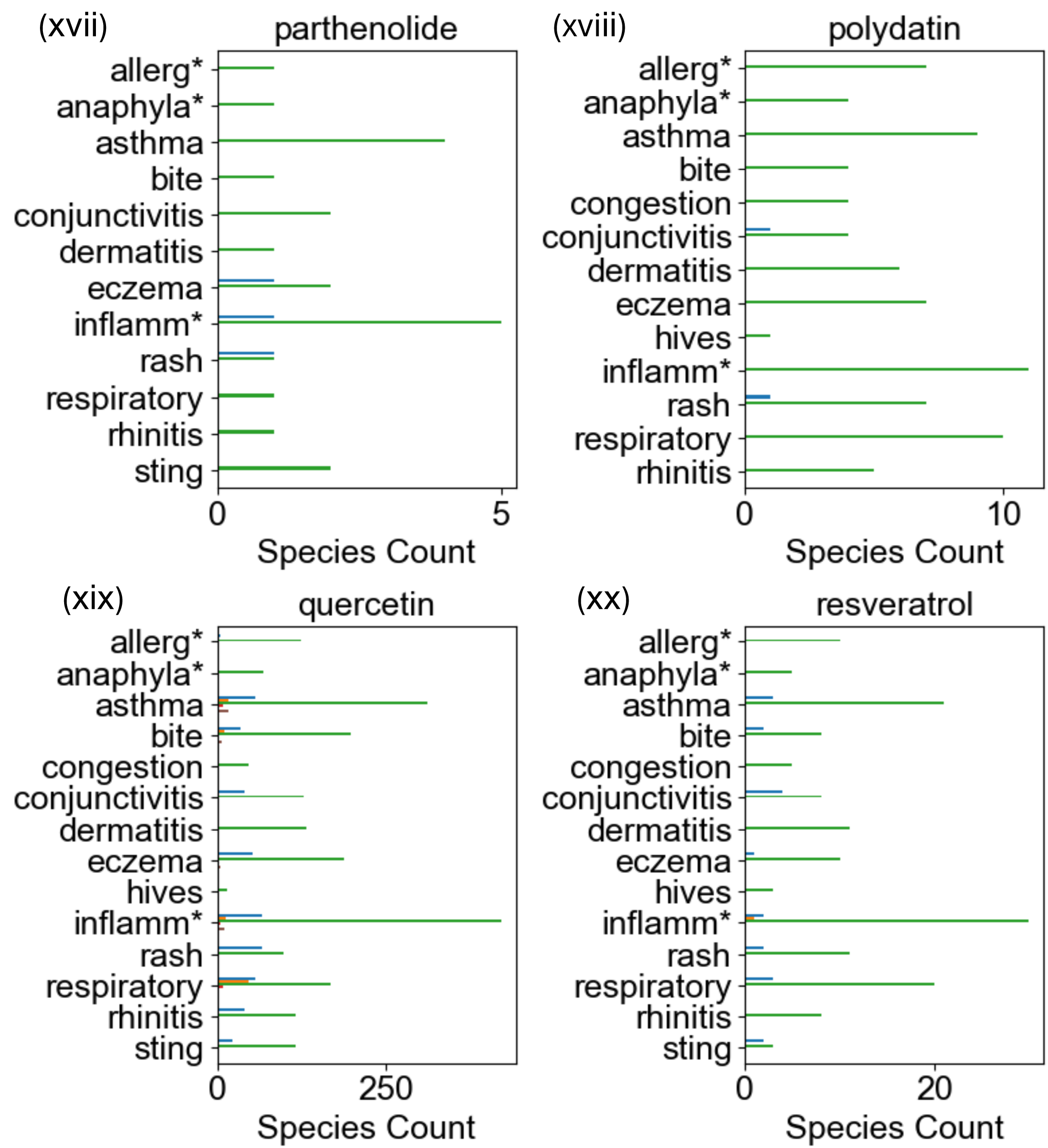
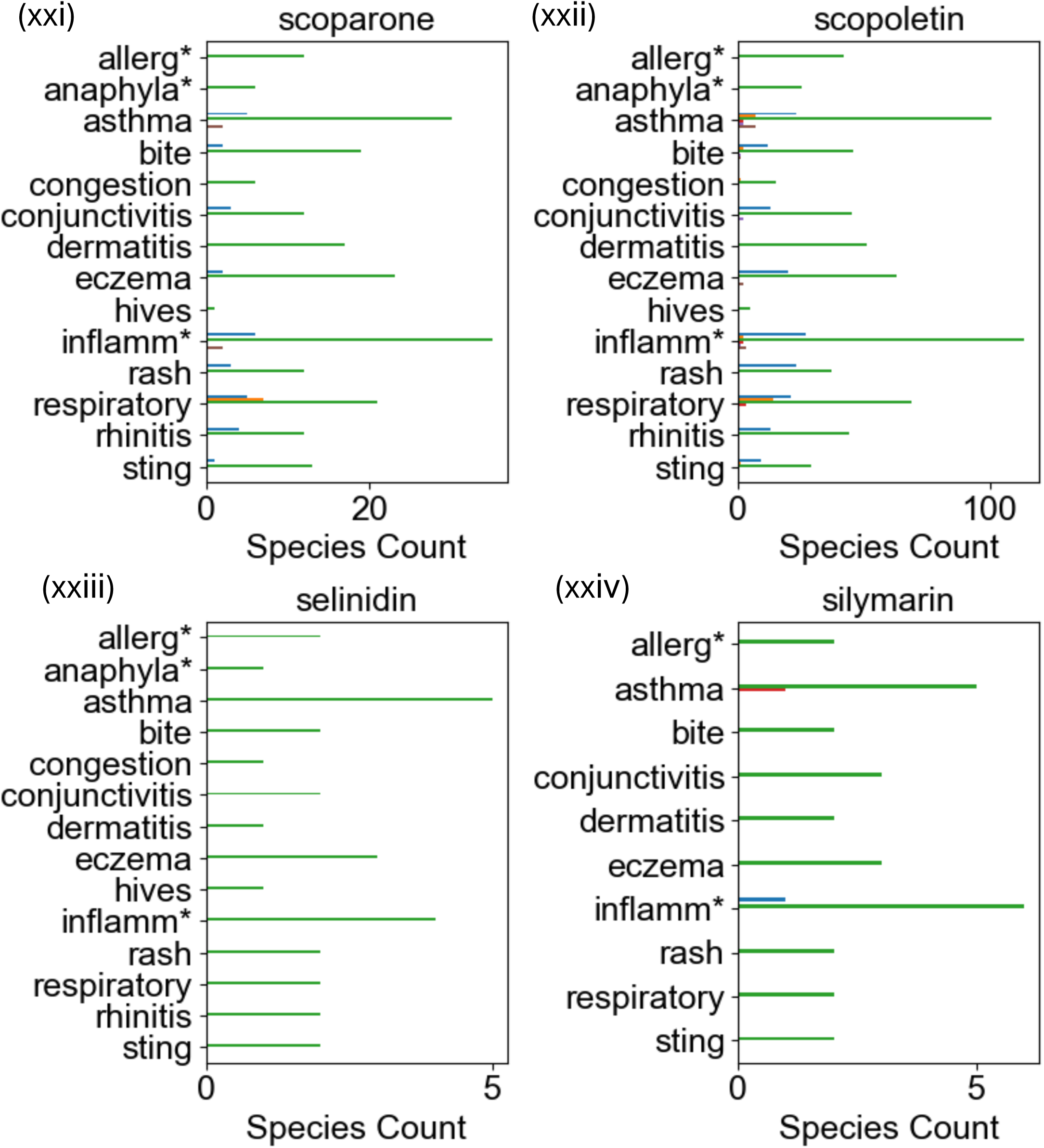
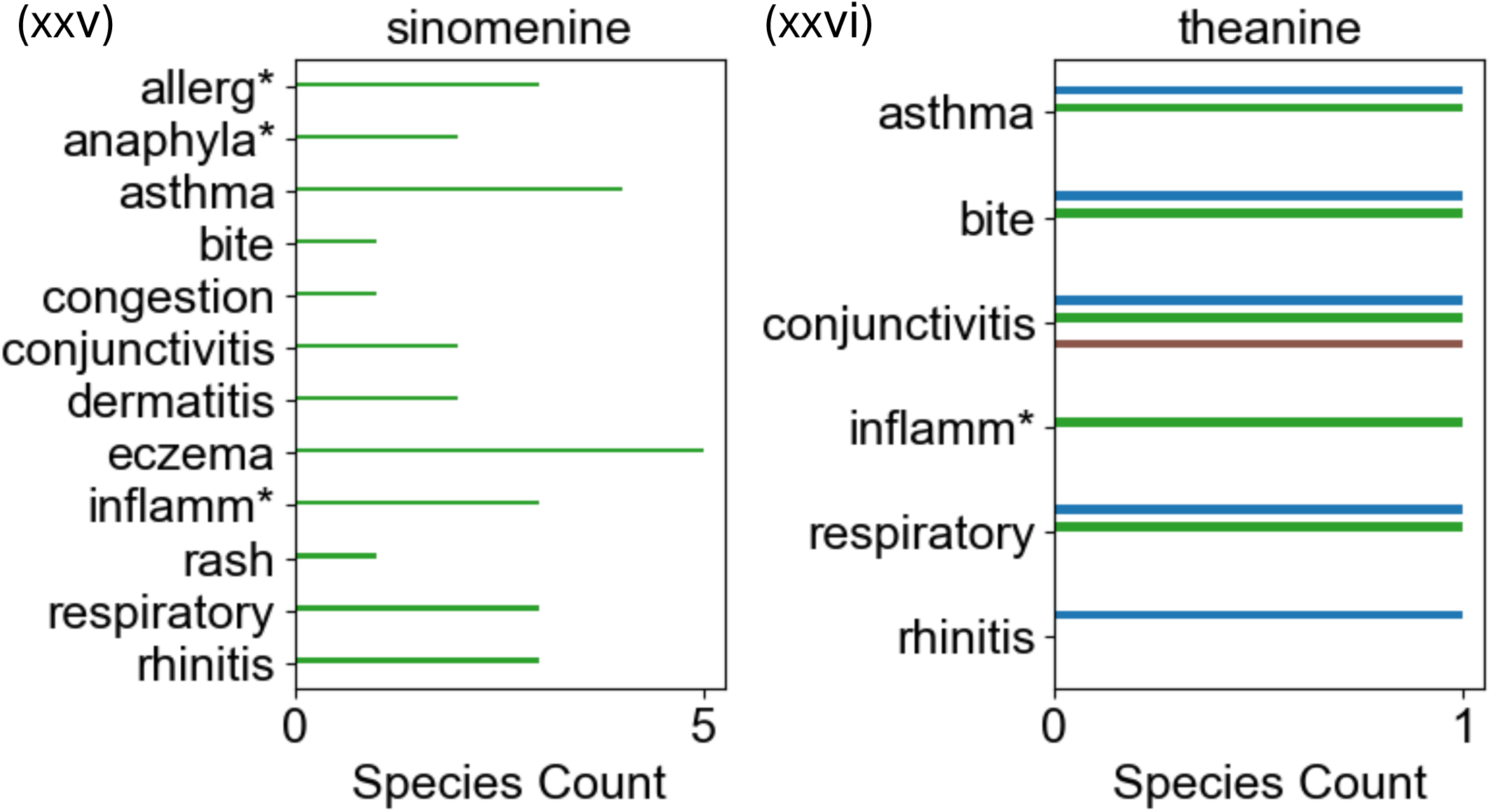
Comprehensive data integration and candidate compound linkage analysis in the Phase I PhAROS^TM^ platform. **A. Overview of the *in silico* convergence analysis (ISCA) environment within the PhAROS^TM^ platform.** Integrating standardized data from eight global traditional medicine systems (IM, AM, EM, CM, JM, KM, SAM) across >15 databases and pharmacopeias. The data include biogeographical, sociopolitical, and temporal contexts (300 BCE–present) for formulations, medical indications, ingredient organisms, and natural compounds. **B.C. Summary of ISCA coverage.** >17,000 formulations, >25,000 ingredient organisms, >75,000 chemical compounds, and >50,000 medical indications sourced from >150 countries and >3,000 cultural groups. Estimated network complexity: >6.9 billion pairwise linkages and 3.1 billion multicomponent paths connecting formulations, indications, organisms, and compounds with other data layers. **D. Candidate ngMCS in PhAROS^TM^**. Heatmap showing linkage of candidate mast cell stabilizers (MCS) from Finn and Walsh’s list to >3,800 instances of ingredient organisms across formulations targeting allergic and inflammatory conditions, based on the constructed indication dictionary (see Methods, Table I). Numbers in parentheses indicate the count of species in PhAROS containing that compound, while the heat map right hand columns represents the % of that count with indications in the specific therapeutic area. **E. Distribution of candidate MCS compounds across global medical systems.** Nineteen of 29 Finn/Walsh compounds are present in at least three systems, indicating widespread utilization. **F. Specific representation of MCS compounds in formulations linked to mast cell-related disorders, disaggregated by system**. Panels (i–xxvi). Notable example: alpha-Mangostin, utilized across two systems (African, Asian) but associated with mast cell-related indications only in Asian medicine.

We developed an indication dictionary (see Methods, Table I) of search terms to link allergic and inflammatory indications for MCS to compounds from the candidate list and the plant species in which they are found across the GMS in the Phase I PhAROS^TM^ platform. We searched the data platform for compounds suggested by Finn and Walsh [20] as potential ngMCS, and cross-referenced them to anti-inflammatory uses for formulations in which they occurred. The heatmap in Figure 2D shows that the candidate MCS compounds links to 3860 instances of an ingredient organism in a formulation associated with one of the indications of interest containing that compound. This suggests that each of the compounds in the Finn/Walsh candidate list are indeed widely utilized in phytomedicinal approaches to mast cell-linked disorders.

We further evaluated the distribution of phytomedicines containing the Finn/Walsh candidate compounds and indicated for mast cell disorders, disaggregating the data by the GMS of origin. Figure 2E shows that 19 of the 29 candidate MCS compounds are represented in at least three GMS suggesting widespread deployment of these compounds in across all indications. We then evaluated each of the candidate MCS for specific representation in GMS formulations associated with the mast cell disorder indication dictionary we constructed (Table I). Figure 2F (panels (i) to (xxvi). Overall, the data supports the Finn/Walsh candidate MCS are represented in traditional/non-Western formulations for one or more of the indicated Mast cell-related disorders. Some specific instances stand out: For example, alpha-Mangostin is a component of phytomedical formulations across two medical systems (Figure 2E, African, Asian) but its deployment in formulations associated with Mast cell related disorders is only seen in Asian medicine (Figure 2F (i)). Some of the MCS candidates are associated with formulations that are used across the indication dictionary, but some are more focused on a limited subset of applications. Mangostins are associated with multiple bioactivities (anti-viral, anti-tumorigenesis) and with effectiveness in experimental models of osteoarthritis and rheumatoid arthritis [66,67]. In the context of the phytomedicine space assessed here, their usage aligns primarily with eczema and we note that Mangostin has been demonstrated clinically to suppress eczema, psoriasis and atopic dermatitis [68–70]. The data in Figure 2F illustrate a first level of decision support capability in the proposed workflow, i.e., de-risking compounds by evidencing their usage in traditional medicine solutions to a disorder of interest and allowing comparison of possible indication profiles to match to unmet therapeutic needs.

### Druggability and target comparison of MCS candidate phytomedical compounds

We curated data on the Finn/Walsh MCS candidates to evaluate indices relating to their relative strength as a natural product drug candidate. Figure 3 (Table II) shows the unique PubChem CID for the Finn-Walsh MCS candidate compounds and we then scored each candidate for several parameters: (1) MW (a range between 259-450 Da is considered desirable); (2) violations of Lipinski’s Rule of Five (Ro5) noting that a compound is likely to have poor absorption or permeation (oral bioavailability) if it violates two or more Lipinski criteria (MW ≤ 500 Da, LogP ≤ 5, Hydrogen bond donors ≤ 5, and Hydrogen bond acceptors ≤ 10); (3) Weighted Quantitative Estimate of Drug-likeness (wQED, 1 = ideal drug likeness, 0 = poor) reflecting how physicochemical properties align with those typically found in successful drugs (including those inherent to Ro5 plus others such as Topological Polar Surface Area and rotatable bond flexibility); and (4) Natural Product Likeness (NP Likeness) where scores from 0–3 are associated with increasing NP characteristics. NP Likeness incorporates structural, physicochemical, and fragment features and its predictive value derives from the assumption that natural products have evolved to interact with biological systems, often exhibiting high specificity and potency. High NP Likeness indicates lead candidate potential. Figure 3 (Table II, A and B) project the QED and NP Likeness druggability indicators across their ranges for all phyto-compounds in the PhaROS data platform. The desirable ranges for the two indicators are shown, allowing both comparisons between the candidates and the total set of phytochemicals. This stage in the workflow again illustrates the decision support capacity of data layers in PhaROS to discern between candidate compounds for follow-up.

**Figure 3.**
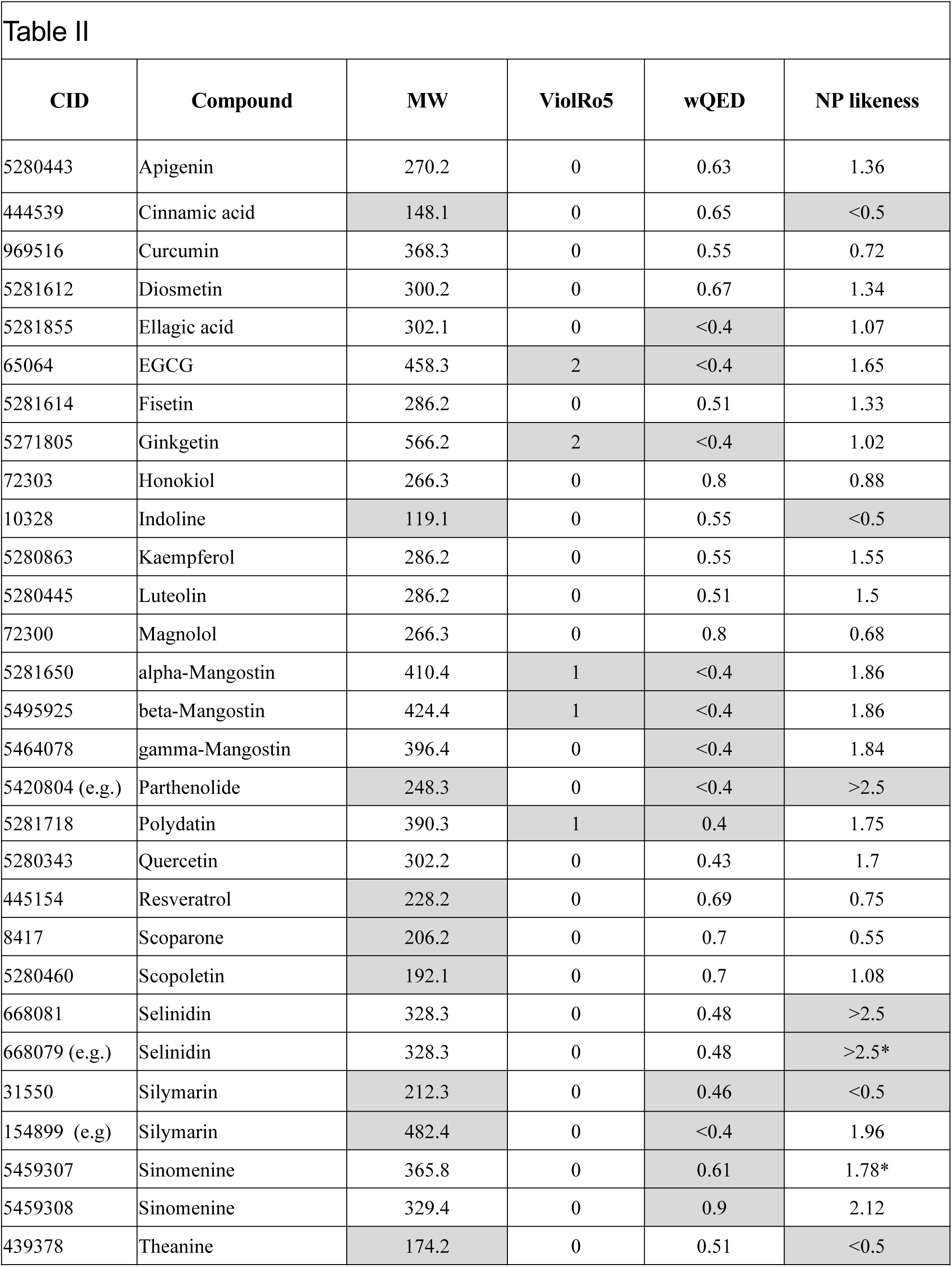
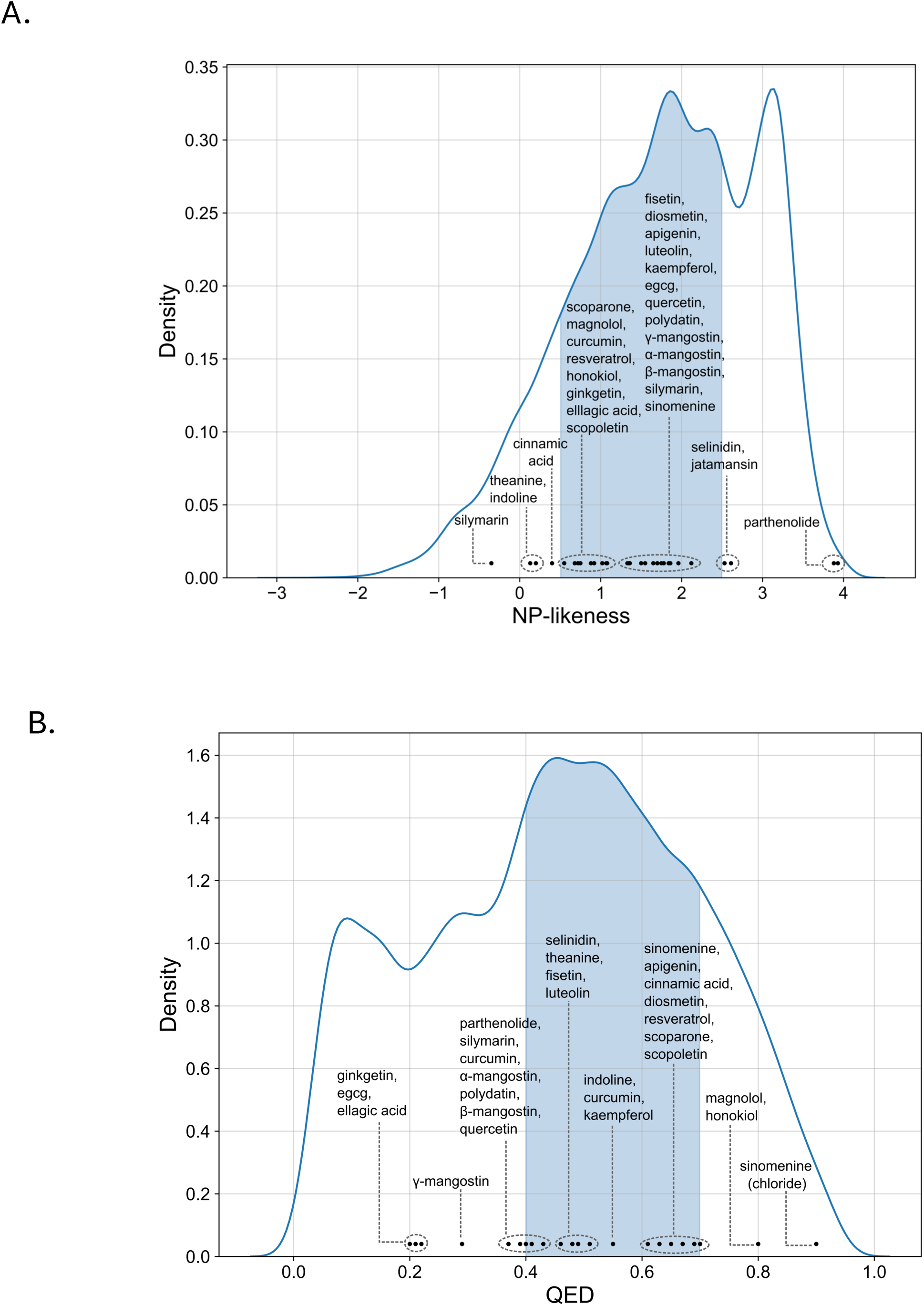
Testing of initial candidate comparison capacity in PhAROS using drug-likeness and chemical characteristics. Table II. Initial discrimination of MCS candidates via drug-likeness evaluation and chemical characteristics. The table presents the unique PubChem Compound Identifier (CID) for each candidate in the Finn/Walsh list, an exemplar SMILES (Simplified Molecular-Input Line-Entry System) representation of its chemical structure, and the corresponding evaluations of key druggability metrics. Each compound is assessed for: (i) violations of Lipinski’s Rule of Five (Ro5), which predicts poor absorption or permeation when two or more criteria are violated, (ii) Weighted Quantitative Estimate of Drug-likeness (wQED), scoring compounds between 0 and 1 based on their alignment with physicochemical properties typical of successful drugs, and (iii) Natural Product (NP) Likeness, an index scoring compounds from 0 to 3, with higher values indicating greater similarity to natural products. Gray shading indicates where a compound ‘fails’ to meet desirable ranges on one of ore of these parameters, lowering its priority for further evaluation. These metrics provide a predictive framework for assessing oral bioavailability, structural drug-likeness, and potential for lead optimization in drug development. **A, B. Projection of QED and NP Likeness druggability indicators for phyto-compounds in the PhAROS^TM^ data platform**. QED (A) and NP-likeness (B) scores for phyto-compounds tabulated across their observed ranges, optimal ranges shaded blue and Finn/Walsh list positions noted.

### Target and mechanism of action analysis in support of compound prioritization

Assessing candidate ngMCS based on their targets and potential mechanisms of action has the potential both to help prioritize candidates and to provide novel insights into how these stabilizers work, given that the mechanisms of action for MCS in current clinical use remain enigmatic. We leveraged data layers in PhAROS^TM^ to map targets for the candidate ngMCS comparing molecular targets between phytochemical ngMCS, and between ngMCS and extant clinical MCS (Tranilast and Chromoglycate). Figure 4A shows a full MCS candidate-target network for the Finn/Walsh compounds, tranilast and cromolyn sodium with dark-blue edges represent validated associations, and light-blue edges represent predicted ones. Figure 4B shows an expanded network visualization including tranilast, cromolyn sodium, and all Finn/Walsh candidate MCS compounds that share at least one known or predicted target with either reference drug, building on the network structure from Figure 4A. Since there is sparse information on the receptors that directly mediate the effects of clinical MCS on mast cells, we filtered the network to focus solely on membrane proteins (ion channels and transporters). This visualization (Figure 4C) is useful for identifying shared potential targets and contains reassuring evidence of targets previously ascribed to MCS (e.g., Chromoglycate and CFTR). It highlights potential mechanisms of action that have previously been proposed for MCS, such as targeting Cl⁻ channels that control the driving force for calcium entry during MC activation or the activating calcium channels themselves. It also brings forward new potential targets in the ABCC and SLC transporter families. We note that there is a high level of representation in the ngMCS candidate group of compounds that target MC calcium influx directly or indirectly, examples including ligands for TRP ion channels that are expressed in mast cells and mediate calcium entry (e.g., alpha mangostin, apigenin, cinnamic acid, curcumin, EGCG, kaempferol, parthenolides, quercetin, resveratrol [71–76]) and natural product regulators of KCa1.3 [77] (e.g., reseveratrol), which modulates the driving force for mast cell calcium entry.

**Figure 4.**
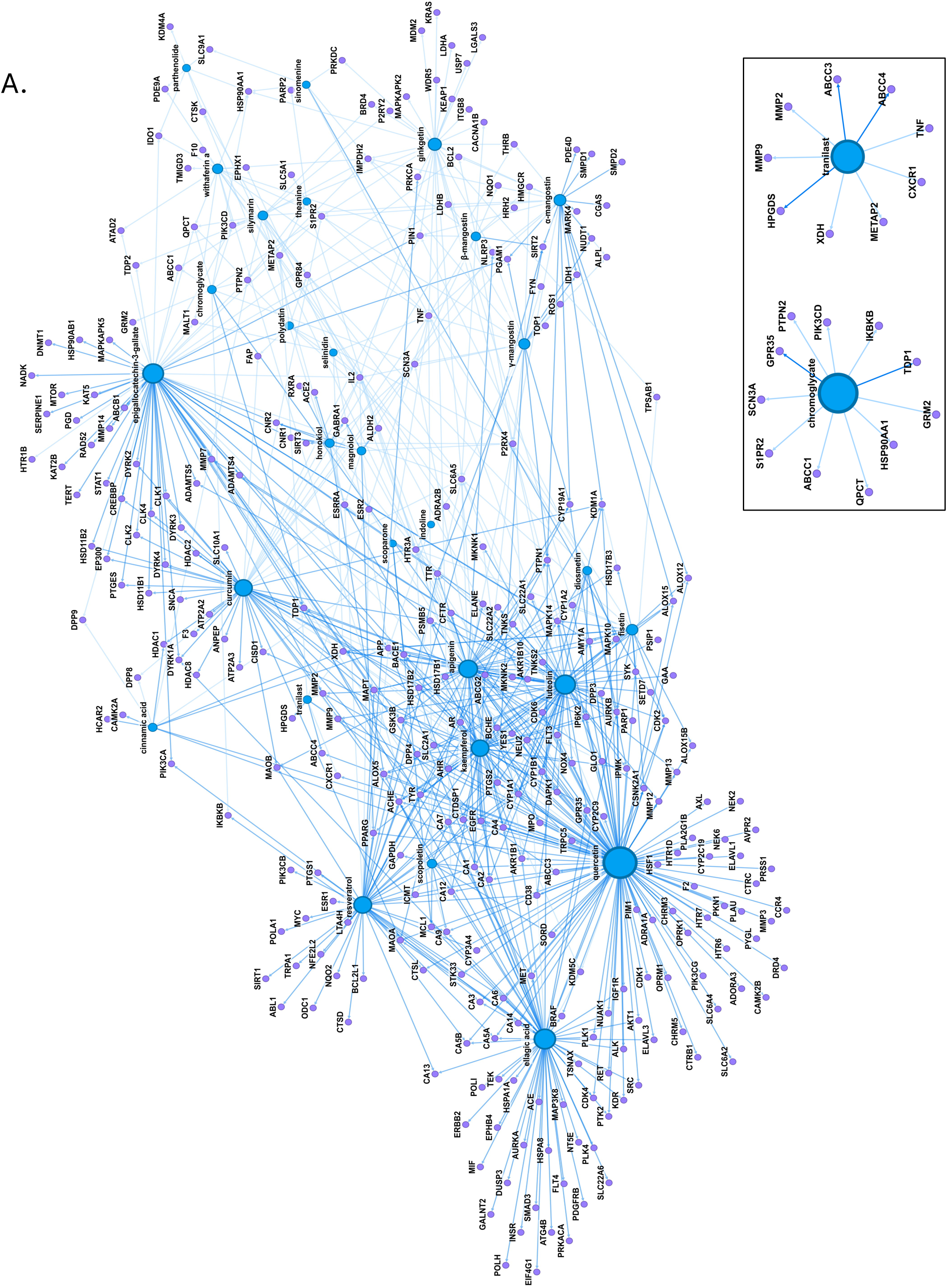
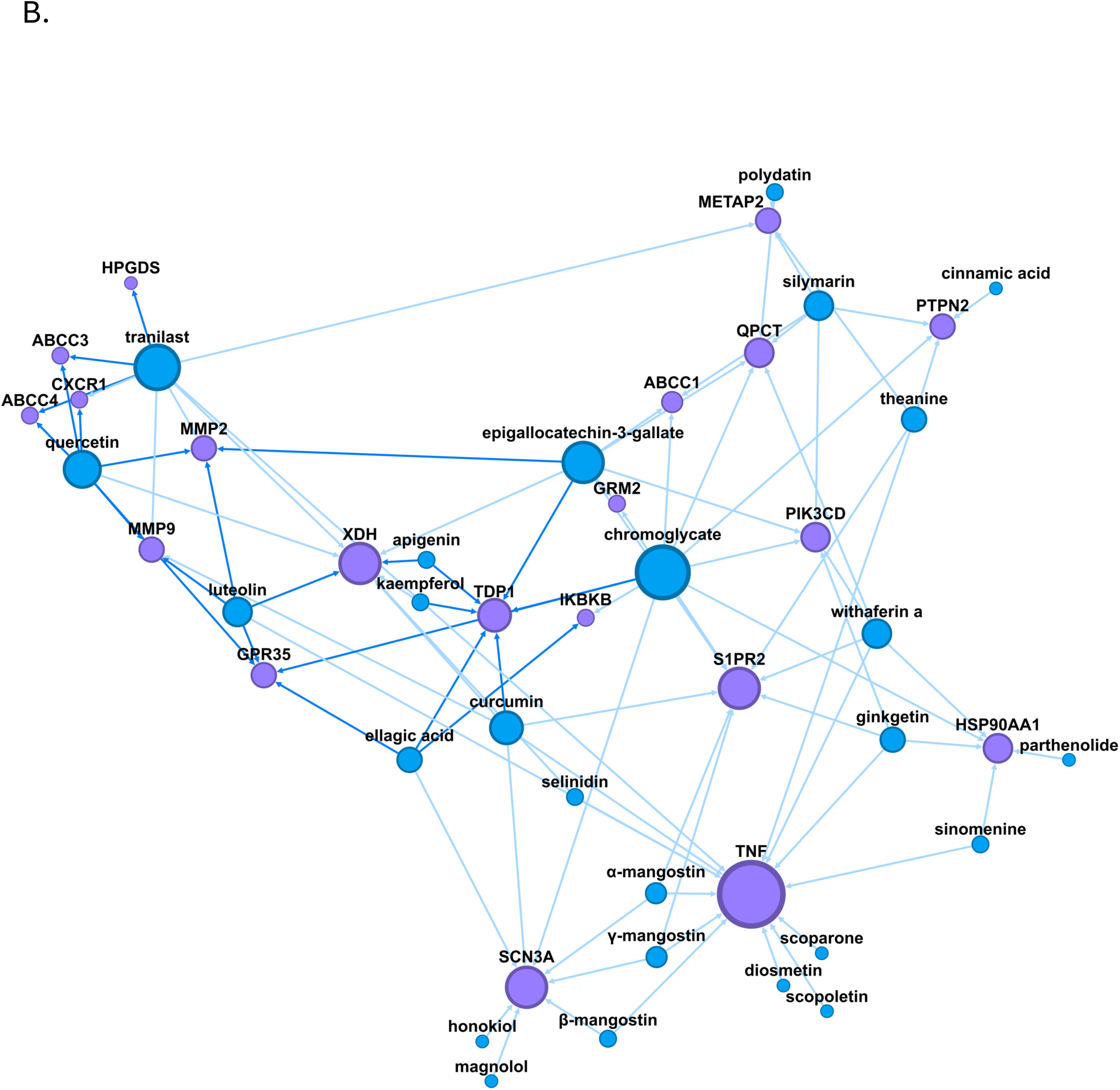
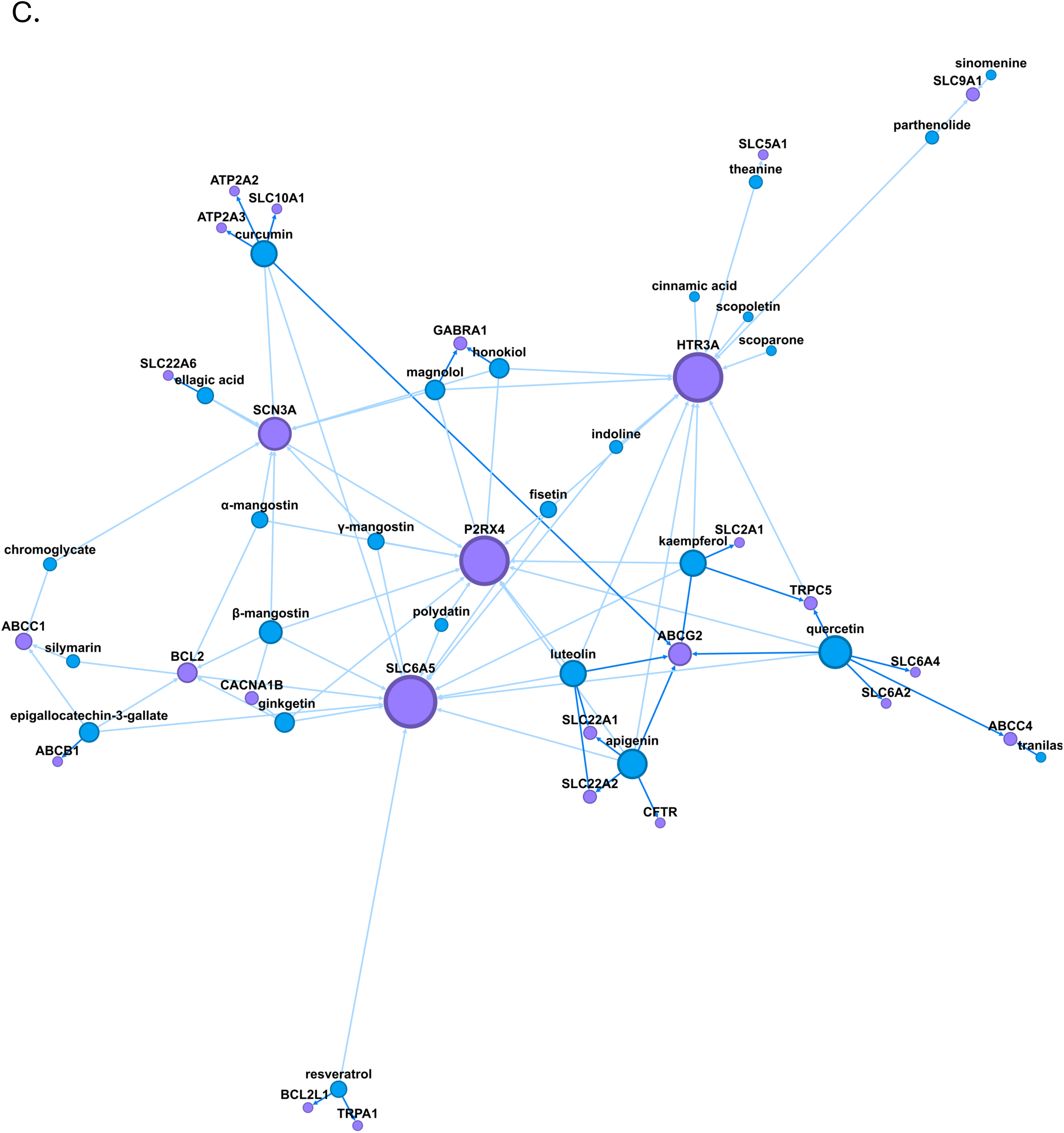
Network plots of ligand-target relationships for clinically established and ngMCS. **A. Full MCS candidate-target network.** *Main panel.* Comprehensive network visualization of ligand-target relationships for tranilast, cromolyn sodium, and all Finn/Walsh candidate MCS compounds. *Inset at right.* Abstracted target networks for Tranilast and cromolyn sodium. Binding associations were sourced from BindingDB and ChEMBL for experimentally validated targets, and from the ChEMBL target prediction API (≥90% confidence, “active”) for predicted interactions. Dark-blue edges represent validated associations, and light-blue edges represent predicted ones. Network data were constructed using Python and NetworkX, and visualized in Gephi using the OpenOrd and ForceAtlas2 layout algorithms. **B. Natural compounds sharing targets with tranilast or cromolyn sodium.** Expanded network visualization including tranilast, cromolyn sodium, and all candidate MCS compounds that share at least one known or predicted target with either reference drug, building on the network structure from Figure 4A. Binding associations were sourced from BindingDB and ChEMBL for experimentally validated targets, and from the ChEMBL target prediction API (≥90% confidence, “active”) for predicted interactions. Dark-blue edges represent validated associations, and light-blue edges represent predicted ones. Network data were constructed using Python and NetworkX, and visualized in Gephi using the OpenOrd and ForceAtlas2 layout algorithms. **C. Ion channel and transporter-focused MCS target network.** Subnetwork derived from Figure 4c, filtered to include only ligand-target associations involving ion channels or transporters—two target classes implicated in mast cell degranulation pathways. Binding associations were sourced from BindingDB and ChEMBL for experimentally validated targets, and from the ChEMBL target prediction API (≥90% confidence, “active”) for predicted interactions. Dark-blue edges represent validated associations, and light-blue edges represent predicted ones. Network data were constructed using Python and NetworkX, and visualized in Gephi using the OpenOrd and ForceAtlas2 layout algorithms. **D. Known and predicted protein targets of tranilast and cromolyn sodium.** Network visualization of compound-to-target relationships between tranilast and cromolyn sodium (blue nodes) and their known or predicted protein targets (purple nodes). Binding associations were sourced from BindingDB and ChEMBL for experimentally validated targets, and from the ChEMBL target prediction API (≥90% confidence, “active”) for computationally inferred interactions. Dark-blue edges represent validated associations, while light-blue edges indicate predicted ones. Network data were constructed using Python and the NetworkX package, and visualized in Gephi using the OpenOrd and ForceAtlas2 layout algorithms.

We note that shortcomings of this network include its limited representation of mast cell calcium channels, which are established targets for multiple candidate ngMCS in the Finn/Walsh list, particularly the terpenes and sesquiterpene lactones. These are mechanistically important because agonist-mediated desensitization [78] of these channels may be a component of the mechanism by which terpenes act as stabilizers (Jansen, data not shown). However, despite there being relevant published studies affirming a relationship between for example terpenes and TRPV1/TRPA1 [78,79], the ontology of the datasets used to generate these networks may place their associations at a lower order (below a certain association threshold where inclusion leads to unmanageable visualizations) or the datasets may simply lack deposited bioassay data. These issues illustrate the need for the workflow in Figure 1 to be augmented with targeted analyses based on specific hypotheses, such as the potential for ngMCS to act via desensitization of MC calcium channels or regulate driving forces for calcium entry via Cl⁻ or K⁺ channels.

### Development of a MCS index score in support of compound prioritization

In the workflow piloted here, we started with a list of candidate ngMCS suggested by Finn and Walsh based on a literature metareview. As an illustrative case study for the use of PhAROS to prioritize these compounds, we followed the schematic in Figure 1 and added various data layers to improve discernment between these phytochemicals, theoretically supporting resource allocation and prioritization for further study. In the final phase of the current study, we interrogated PhAROS^TM^ for an ngMCS candidate list independent of the Finn and Walsh study, but continuing to ground-truth the PhAROS^TM^ outputs with reference to the Finn/Walsh list.

We developed a novel ‘MCS score’ to rank phytochemicals in PhAROS^TM^ based on their likelihood of MCS potential. The initial Harmonic Mean-based MCS Score (HM-MCS) integrates two key parameters: *breadth* (the number of distinct indications of interest associated with a compound) and *depth* (the number of plants containing the compound within the database metapharmacopeia). The harmonic mean was selected because it emphasizes smaller values, ensuring that a high score requires meaningful contributions from both breadth and depth. This prevents overranking compounds with disproportionate profiles-such as those widely cited across indications but rare in phytochemical sources, or those abundant in many plants but linked to few relevant indications. This scoring strategy aligns with the core PhAROS^TM^ design principle of leveraging consensus and convergence across medical systems and formulations to de-risk and prioritize compounds for further study. The HM-MCS thus provides a balanced composite score that penalizes extreme outliers while rewarding consistent, reliable representation across formulations. Figure 5A shows the distribution of the HM-MCS Score across the 36,117 compounds within the indication dictionary-filtered subset, with the Finn-Walsh candidate set annotated. Based on this Tukey plot of HM-MCS scores across all compounds in the set we noted a possible cut-off threshold (top 12% of HM scores). We evaluated the Finn and Walsh compounds using this new HM-MCS Score and noted that of 26 candidate compounds from that study that are represented in PhAROS^TM^, 23 exceeded the mean HM-MCS and 19 exceeded a nominal outlier threshold of top 12^th^ percentile for MCS Score (Figure 5A). We noted (Figure 5B) that this threshold did not specifically disadvantage relevant compound chemical types represented in the data platform. Finally, we approached PhAROS^TM^ independently of the Finn and Walsh candidate MCS list and interrogated the database for the highest HM-MCS scoring compounds. The bubble plots in Figure 5C show that as the positivity of the HM-MCS score increases, the relative compound class representation in the population of candidates also changes. While the cutoffs here are arbitrary (top 1%, 5% and 10% of HM-MCS scores) at this time, they could in the future be associated with preclinical results, allowing us to refine the use of HM-MCS performance as a means to identify classes of chemical with more or less likely potential as ngMCS.

**Figure 5.**
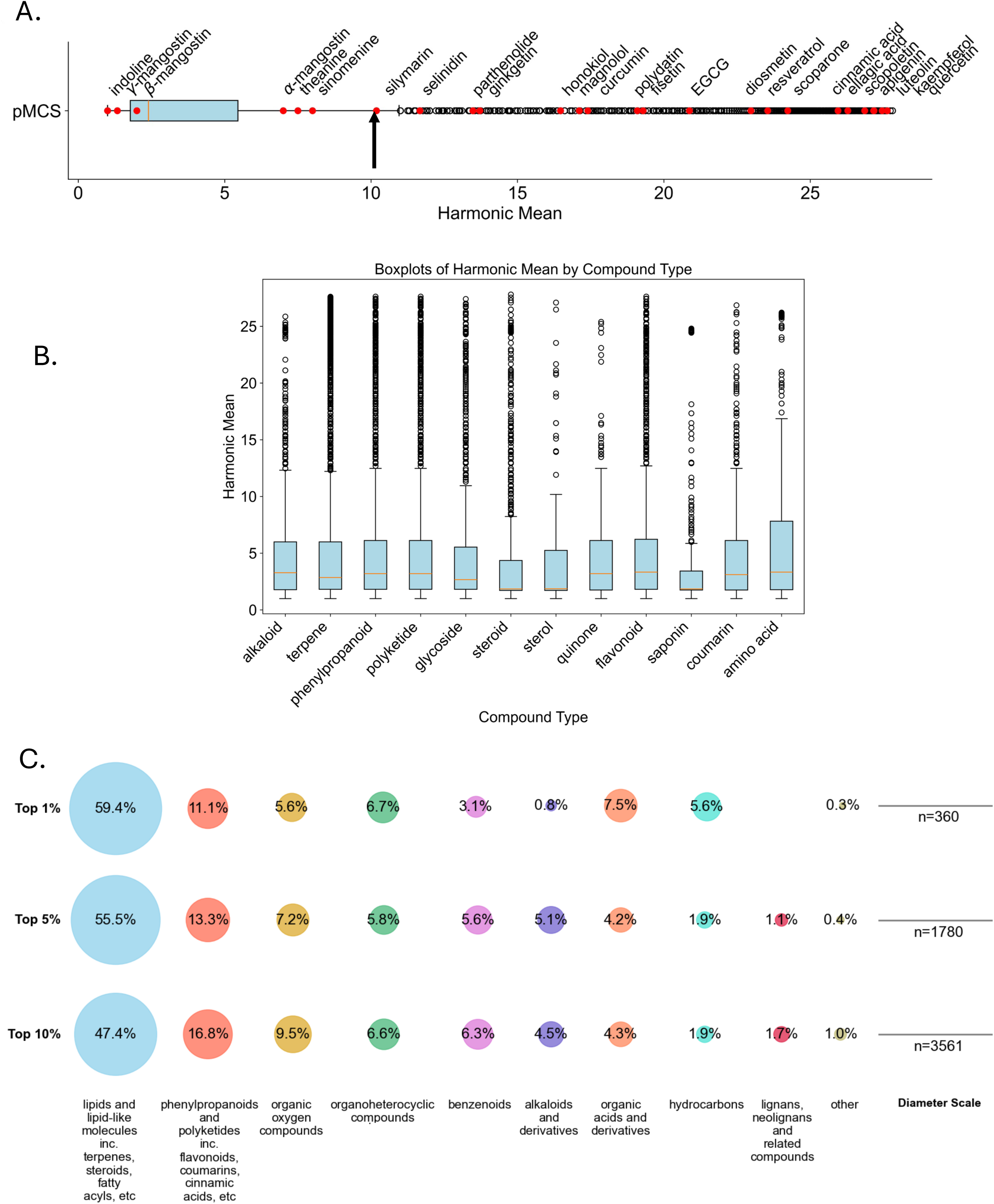
Prioritization and evaluation of candidate mast cell stabilizers (MCS) using the PhAROS Harmonic Mean MCS (HM-MCS) Score. **A. Distribution of the HM-MCS scores for 36,117 compounds filtered from the PhAROS indication dictionary.** Scores are calculated as the harmonic mean of Breadth (number of unique target indications associated with a compound) and Depth (frequency of occurrence across plant species). The Tukey boxplot displays HM-MCS score distribution. The upper outlier threshold (defined as Q3 + 1.5×IQR) is used as a nominal cutoff for prioritization, corresponding to the top 13th percentile. Of 27 Finn and Walsh candidate compounds (red dots), 19 exceed the 12th percentile HM-MCS threshold. **B.** Chemical class composition analysis shows that applying this threshold does not systematically exclude specific compound types. **C.** Bubble plots visualizing HM-MCS scores of compounds independently identified from the PhAROS database, highlighting key candidates with high prioritization potential. This composite scoring system provides an evidence-based framework for de-risking and prioritizing compounds for further experimental validation.

#### Testing of extended HMS indices to rank and prioritize ngMCS candidates in PhAROS^TM^

We tested variations on the HM-MCS scoring index in order to fully leverage the data layers platformed in PhAROS^TM^ and generate refined candidate lists for further investigation. The HMS score (Index I) was treated as a foundational list based solely on traditional use patterns. prioritizing compounds using a harmonic mean of two ethnopharmacological parameters: the number of mast cell–relevant indications (such as inflammation, pain, and allergy) and the number of distinct species in which the compound appeared. This provided a weighted metric favoring compounds that were both widely used across traditional systems and linked to therapeutically aligned applications. Index variation II expanded this approach, introducing cheminformatic prioritization by incorporating biological target overlap with tranilast and cromolyn using shared ChEMBL protein targets, Additionally, we factored in the total number of known ChEMBL targets for each compound, alongside desirable qualities such as high QED, high NP, and low violations of Lipinski’s RO5. This multi-criteria sort aimed to balance traditional utility with drug development potential. However, it is limited inherently by the sparse understanding of the mechanisms and targets for these two clinically-relevant MCS. In Index variation III, we introduced a binary composite score based on key physiochemical parameters. Compounds received one point each for RO5 compliance (i.e., no violations), molecular weight between 200–450 g/mol, QED score between 0.4–0.7, and NP-likeness between 0.5–2.5. The resulting composite score was then multiplied by the harmonic mean score from Index I to prioritize compounds that met both drug-like and ethnobotanical criteria. Index variation IV expanded this approach by including a binary score for compounds with any ChEMBL protein target relevant to mast cell stabilization including histamine signaling, arachidonic acid metabolism, i cytokine pathways, and other signal transduction pathways, each validated by literature search. Finally, Index variation V refined the target-based approach of Index IV by replacing the binary target score with a scaled value based on the number of relevant ChEMBL targets. A logarithmic scale was used to weight target count, ranging from 0 (no relevant targets) to 3 (20 or more targets), with intermediate values scaled appropriately (e.g., 0.7 for one target, 1.8 for five, 2.4 for ten). This produced a continuous-valued target score that was then combined with the existing composite framework and multiplied by the harmonic mean, further elevating compounds with both structural desirability and substantial biological relevance. With these five indexing methods in hand, we ‘ground truthed’ them with the Finn/Walsh candidate set members that exceeded our nominal Index I cutoff (>12^th^ percentile) (Figure 6A), noting that progression from I to V was associated with increasing performance of the Finn/Walsh compounds.

**Figure 6.**
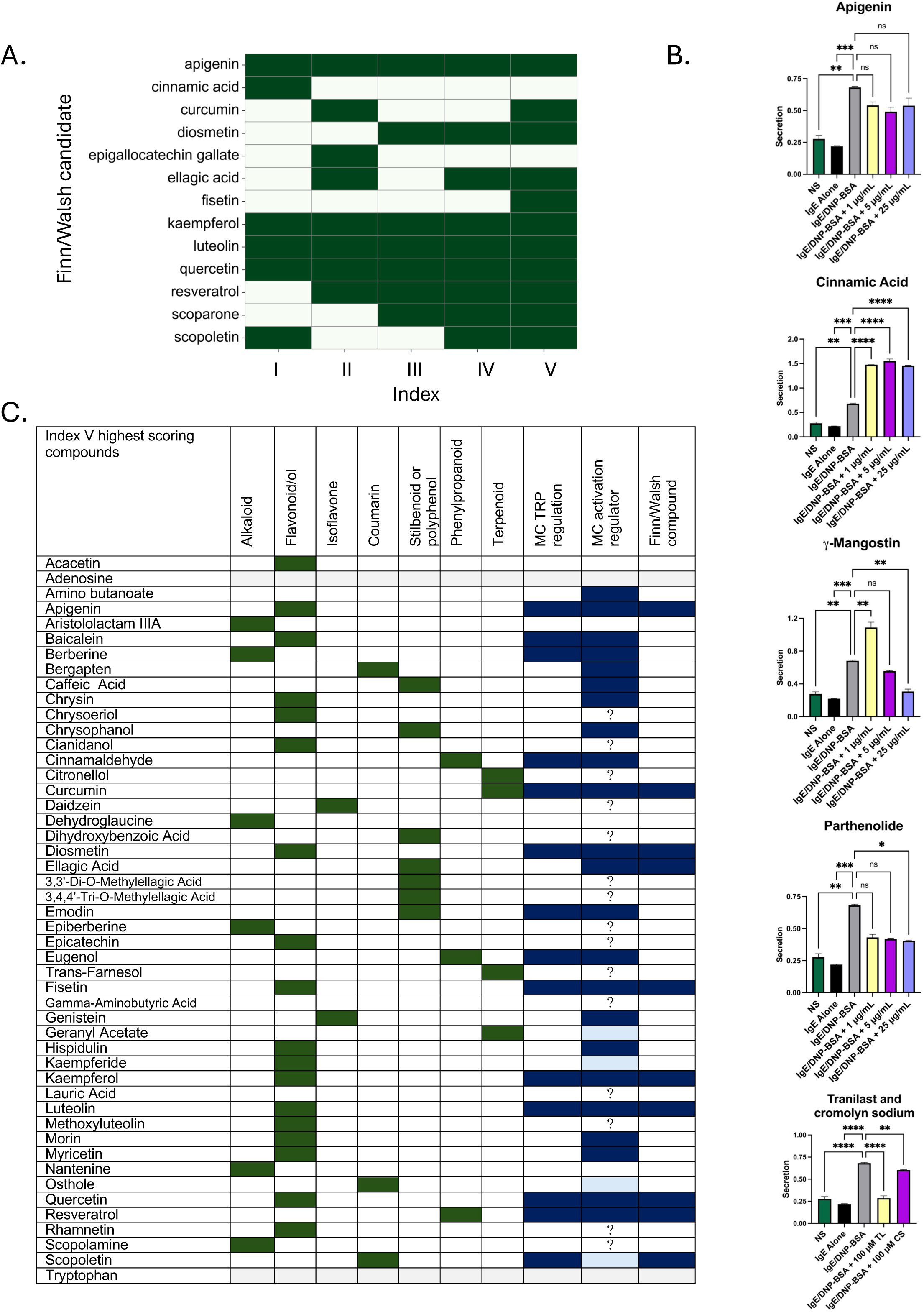
Extended novel indices for ranking candidate ngMCS. **A. Heatmap of Finn and Walsh benchmark compound presence across compound ranking methods.** The y-axis lists benchmark compounds derived from Finn and Walsh that met initial selection criteria based on a harmonic mean of ethnopharmacological relevance and traditional usage breadth. The x-axis represents extended ranking methods (Index I through Index V), each incorporating increasingly refined criteria including drug-likeness and molecular target relevance (see *Methods*). A green-filled box indicates that the compound was ranked among the top 100 by the respective method; blank boxes denote absence. This heatmap visualizes how each scoring strategy recapitulates or omits known or suspected mast cell stabilizers. **B. Candidate MCS were evaluated in MC degranulation assay**. Degranulation was assessed using β-hexosaminidase release, assay following stimulation via the high-affinity IgE receptor, FcεRI. **C. Index V highest scoring candidates in PhAROS^TM^.** highlighted by documented TRPV ion channel regulation, documented mast cell regulatory effects, and chemical class [33,73,83–116].

We performed in vitro testing (Figure 1) on three Finn/Walsh compounds that illustrate three scenario outcomes of the HM-MCS Indexing, using a mast cell degranulation bioassay (Figure 6B). Apigenin performs well across all V indexing approaches, and is a validated TRV1 ligand with the potential to stabilize mast cells through agonist mediated desensitization of calcium entry pathways. It tends to suppress degranulation in our assays. Cinnamic acid performs moderately in the HM-MCS indexing but is primarily a TRPA1 ligand (a channel also expressed in MC and mediating calcium entry). It actually stimulates degranulation (pro-inflammatory), suggesting that discriminating between TRP channel targets would be important step going forward in prioritizing candidates. Mangostins and Parthenolide perform at a lower level in HM-MCS, with only one species exceeding the mean of all PhAROS^TM^ compounds. They are effective in a mast cell activation bioassay but their relatively poor performance in rankings is driven by druggability issues (Table II in Figure 3), which could suggest they be targeted for bioavailability improvements [80–82]. This reinforces the component in our workflow (Figure 1) of a ‘human-in-the-loop’ hypothesis testing component in the candidate ranking and prioritization process. We also tested both, cromolyn sodium and Tranilast, known MCS, and saw suppressed degranulation.

Finally, we used PhAROS^TM^ as a predictive tool to generate a *de novo* prospective ngMCS set comprising Index V ∼50 highest scoring compounds (Figure 6C). This list contains some Finn/Walsh compounds but extends beyond them. The chemical classes most associated in PhAROS^TM^ with MC-related indications predominate in the highest scoring Index V candidate list and after generating generic/limited bioactivity components (e.g., adenosine, tryptophan), then 53% of high-performing compounds have some documented MC regulatory history and/or capacity to regulate MC calcium entry as a mechanism of action [33,73,83–116]. Several are positioned for further exploration as they have not previously been associated with MC therapeutic potential, and some would be potentially placed at a lower priority level based on their compound class and lower indexed score ranking.

## Discussion

Mast cell stabilizers (MCS) appear to operate on multiple levels beyond the conventional focus of antihistamines, corticosteroids, and biologics, potentially offering long-term control in allergic and inflammatory conditions such as mast cell activation syndrome (MCAS), atopic dermatitis, and asthma. By targeting multiple aspects of mast cell mediator release and exerting chronic stabilizing effects, ngMCS could address some gaps in current therapeutic strategies. However, the clinical use of MCS has been limited by issues such as poor bioavailability, frequent dosing requirements, and an unclear mechanism of action, which have constrained their integration into standard treatments for allergic and inflammatory disorders. Our study implemented a data-driven strategy that integrates global pharmacopeias, data science, and preclinical pharmacology to streamline ngMCS discovery from phytomedicinal sources.

The PhAROS^TM^ data platform and analytical pipeline links numerous layers of data across historical and contemporary non-Western medical systems, enabling a systematic mapping of phytomedicine-derived compounds to validate mast cell-related indications. The results suggest that candidate ngMCS exhibit conservation across multiple traditional medicine systems, reinforcing their historical use as mast cell-targeted therapies. The inclusion of druggability indices in our analysis further supports the prioritization of candidates based on physicochemical properties, structural compatibility with known drug-like compounds, and bioavailability potential. We note, however, that bioavailability *in situ* in phytomedicines is likely influenced by polypharmacy, including co-formulation with excipients and encapsulants. In parallel work (in preparation) we have started to incorporate a heuristic of ‘rules of assembly’ based on the Chinese 君臣佐使 jūn chén zuǒ shǐ) system [117], where components in phytomedicines are classed as ‘Chiefs, Deputies, Assistants and Envoys’ (CDAE). This work allows us to optimally design and pair multicomponent mixtures for wet lab testing based on PhAROS^TM^ outputs. Bioavailability remains a key limitation of current mast cell stabilizers; addressing this challenge should be a priority in determining which candidate compounds to advance for further development. The CDAE work we are performing is allowing, for example, us to identify relevant candidate excipients, encapsulants and adjuvants (E and A components) to be paired with the bioactive primary (C) compounds we identify in studies like the one presented here. Our proposal of the harmonic mean-based MCS score offers an example of a quantitative metric for prioritizing compounds. By balancing breadth (representation across different medical systems) and depth (frequency within formulations), this score facilitates compound ranking and decisions about progressing compounds for future testing. Notably, 19 out of 29 candidates identified in the Finn and Walsh metareview exceeded our threshold for prioritization, suggesting alignment between phytomedical use and computational predictions. The extension of HM-MCS scores into 4 additional indices that differentially balance ethnopharmacological insights with druggability and target prediction has laid a foundation for Machine Learning. In ongoing studies, we are now deploying Random Forest models with categorical booting and SHAP to identify and rank importance features that drive Index I-V scores as a dependent variable. This next stage of the project will fully realize PhAROS^TM^ as a predictive analytics tool. In the current study, interrogation of PhAROS^TM^ on the basis of HM-MCS indices, independent of the starting Finn/Walsh candidate list, yielded a candidate list that included some Finn/Walsh compounds, compounds that fall outside that set but with documented promise as MC regulators, and compounds that have not been explored experimentally for MC regulation but are highly prevalent in ethnomedical solutions to MC-driven disorders [65].

We note that Khellin (the African/Middle eastern phytomedical compound that was the original source for cromoglycate) performed only moderately well (HM 6.66) in the HM-MCS score. This reflects its limited breadth (few plant species contain it, they have limited biogeography and they are not highly represented in PhAROS^TM^). Khellin is likely to act through GPCR, as its regulation of adenylyl cyclase in a G αi/o dependent fashion has been documented [118,119]. However, its direct molecular targets are not resolved, suggesting that progression through indices II-V, with their incorporation of target binding and bioassay data, would not improve its ranking dramatically. This suggests HM-MCS is not a stand-alone measure and perhaps underscores the need for ML to identify a broad range of chemical, biochemical and ethnopharmacological importance features that can then be used to rank and evaluate compounds into candidate sets for a specific indication. Additionally, the origin plant for Khellin identified in the Altounyan studies (*Ammi visnaga)* is present in PhAROS and associated with African, Asian, European and American medical systems. A strength of PhAROS is that it tells us that Khellin is also found in additional species *(Ammi majus*, *Dioscorea deltoidea*, *Dioscorea parviflora*, *Trillium camtschaticus)* which can now be assessed for inclusion formulations that address Mast cell -associated disorders and assessed for potential polypharmaceutical pairings using the CDAE heuristic described above.

There is a need for further mechanistic exploration of ngMCS to better understand their precise molecular targets and pathways, as current clinical MCS lack well-defined mechanisms of action. Our comparative target analysis using ligand-target network mapping provided some initial insights into potential molecular mechanisms of action for ngMCS candidates. Our findings reaffirm previously proposed mechanisms involving chloride channels (CFTR) and calcium channels while also identifying novel potential targets within the ABCC and SLC transporter families. The striking representation of TRP ion channel ligands in both the Finn/Walsh candidate set and our predictive data are intriguing because agonist-mediated desensitization (AMD) of these ion channels is a potential mechanism of action hypothesis for MCS that our group is actively investigating. The need for coupled bioassay work in addition to data science is underscored by the example of the distinctions between the effects of apigenin which targets TRPV1 and Cinnamic acid which targets TRPA1. Both channels mediate MC calcium entry and both compounds are highly associated with anti-inflammatory phytomedicine formulations across medical systems. However, the *in vitro* bioassay integration in our workflow (Figure 1) let us distinguish between them for further exploration and informed our understanding of the differential roles of ion channel players in the AMD mechanism.

While this study offers a structured approach to identifying and prioritizing ngMCS, several limitations are acknowledged. First, the complexity of global medicine formulations poses challenges in isolating individual bioactive compounds. Many phytomedicine-derived treatments consist of mixtures of compounds that may exert interacting effects, making it difficult to determine the specific contribution of each component. Future studies should incorporate polypharmaceutical and synergy assessments to account for these interactions. Second, the lack of standardized bioavailability and pharmacokinetic data for many phytocompounds necessitates further optimization efforts, including formulation improvements to enhance stability and systemic absorption (including those suggested by the CDAE framework described above). Third, while in vitro assays provide critical early-stage validation, they do not fully capture the complexity of mast cell biology in a physiological context. In vivo studies are essential to confirm efficacy, pharmacokinetics, and potential off-target effects. Additionally, all computational predictions and data-driven prioritization strategies, while useful, are inherently limited by the quality and completeness of the underlying data. Phytochemical databases and global medicine compendia may contain inconsistencies, and the predictive power of computational models is constrained by available bioassay and target data.

This study presents an integrative approach to identifying and prioritizing ngMCS derived from phytomedicine using a combination of computational analytics and in vitro validation. Our findings provide evidence supporting the historical and contemporary use of phytocompounds as mast cell stabilizers, suggesting their potential as future therapeutic candidates. The development of the PhAROS data platform and the introduction of the harmonic mean-based MCS score represent methodological advancements that may aid in drug discovery. However, further research is needed to refine mechanistic insights, improve bioavailability, and advance preclinical validation to facilitate the transition of ngMCS from phytomedicine to therapeutic applications.

## Author Contributions (CRediT Taxonomy)

Conceptualization: HT, AJS, CJ, CNA, ALSH. Data curation: BR, BW. Funding acquisition: HT, ALS, ALSH. Investigation: BR, BW, CJ, SE, LS. Methodology, BR, BW. Software: BR, BW. Visualization, BR, BW, CJ. Writing – original draft: HT, CJ. Writing – review & editing: HT, CJ, BR, BW, AJS, ALSH

## Acknowledgements.

The authors thank Ms. Meghan Matsuoka (Undergraduate Program in Biology, Chaminade University), Ms. Julia Howard (Graduate Program in Molecular Biosciences and Bioengineering), University of Hawai’i for research assistance, and Dr. JD Baker (Chaminade University) and Dr. Apo Aporosa (University of Waikato) for helpful discussions.

## Statement on use of Generative AI

Declaration of generative AI and AI-assisted technologies in the writing process during the preparation of this work the author(s) used iChatGPT4o in order to research, edit and check text. After using this tool/service, the author(s) reviewed and edited the content as needed and take(s) full responsibility for the content of the publication.

